# Downsizing in plants - UV induces pronounced morphological changes in cucumber in the absence of stress

**DOI:** 10.1101/2021.02.27.432481

**Authors:** Minjie Qian, Eva Rosenqvist, Els Prinsen, Frauke Pescheck, Ann-Marie Flygare, Irina Kalbina, Marcel A.K. Jansen, Åke Strid

## Abstract

Ultraviolet (UV)-A- or UV-B-enrichment of growth light resulted in a stocky cucumber (*Cucumis sativus* L.) phenotype exhibiting decreased stem and petiole lengths and leaf area. Effects were larger in plants grown in UV-B- than in UV-A-enriched light. In plants grown in UV-A-enriched light, decreases in stem and petiole lengths were similar independently of tissue age. In the presence of UV-B radiation, stems and petioles were progressively shorter the younger the tissue. Also, plants grown under UV-A-enriched light significantly reallocated photosynthate from shoot to root and also had thicker leaves with decreased specific leaf area. Our data therefore imply different morphological plant regulatory mechanisms under UV-A and UV-B radiation. There was no evidence of stress in the UV-exposed plants, neither in photosynthetic parameters, total chlorophyll content, nor in accumulation of damaged DNA (cyclobutane pyrimidine dimers). The ABA content of the plants also was consistent with non-stress conditions. Parameters such as total leaf antioxidant activity, leaf adaxial epidermal flavonol content and foliar total UV-absorbing pigment levels revealed successful UV acclimation of the plants. Thus, the stocky UV-phenotype was displayed by healthy cucumber plants, implying a strong morphological response and regulatory adjustment as part of UV acclimation processes involving UV-A and/or UV-B photoreceptors.

## 1. Introduction

The study of plant UV-responses has gradually shifted from plant stress biology in to the realm of plant regulatory responses (Jansen & Bornman, 2012). There is now good, emerging understanding of the mechanism underlying the sensing of UV-B and UV-A wavelengths by a range of dedicated plant photoreceptors, including the phototropins, cryptochromes and UVR8 (Paik & Huq, 2019). However, understanding of downstream regulatory interactions which can substantially modify UV-A and UV-B responses is only slowly emerging. Nevertheless, there is consensus that both UV-B and UV-A signalling pathways are closely interacting (Rai et al., 2019; 2020), with further crosstalk with, amongst others, phytochrome signalling. For example, UV-B through UVR8, accelerates degradation of PHYTOCHROME INTERACTING FACTORS (PIFs) that are part of the phytochrome mediated elongation response to high far-red to red light ratio’s (Sharma et al., 2019). As a consequence of these interactions, plant responses under natural light conditions are not always identical to those observed under controlled, artificial lighting in the laboratory. For example, Morales et al. (2013) showed that under natural, sunlight, UVR8 both positively and negatively affects UV-A-regulated gene expression and metabolite accumulation. Conversely, a high UV-A/blue light background radiation moderates UV-B-driven gene-expression.

Understanding plant UV-responses under natural conditions is particularly important in the context of climate change. Ongoing changes in the global climate, recovery of the stratospheric ozone layer, and interactions between these two processes, are resulting in novel combinations of, amongst others, temperature, water-availability, and solar UV radiation (Bornman et al., 2019). For example, plants in the Mediterranean are predicted to be exposed to higher UV-levels, due to climate change-associated changes in cloud cover, together with increased spells of drought (Bornman et al., 2019). It has been hypothesised that UV-exposed plants will be more drought-protected. Indeed, Robson, Hartikainen & Aphalo (2015a) showed that when silver birch seedlings were exposed to a combination of natural UV and drought, wilting was less pronounced compared to that in plants which had just been exposed to drought. However, not all studies show such cross-tolerance (Rodríguez-Calzada et al., 2019), and there is still considerable uncertainty in the literature concerning the environmental relevance of cross-tolerance (Jansen et al., 2019). Nevertheless, it has been argued that a key component of any putative cross-tolerance is the UV-induced change in plant architecture, and especially a more stocky phenotype. The UV-induced phenotype is characterised by shorter stems, internodes, and petioles, and a diminished leaf area, often associated with an increase in leaf thickness (Jansen, Gaba & Greenberg, 1998; Robson, Klem, Urban & Jansen, 2015b), and some of these characteristics are shared with drought-acclimated plants. Yet, major questions remain concerning the stocky UV-phenotype, and particularly the mechanism underlying the induction of such a phenotype. UV-induced stress, possibly involving reactive oxygen species (ROS; Hideg, Jansen & Strid, 2013) and/or accumulation of damaged DNA (Kang, Hidema & Kumagai, 1998), may affect plant architecture (Robson et al., 2015b). Conversely, a regulatory response mediated by a UV photoreceptor can drive architectural change. In the latter case, UV-B- and UV-A-induced responses may be different as these are driven by distinct photoreceptors.

The aim of the current study was to investigate whether morphological changes occurred in plants as a result of supplementing photosynthetically active radiation (PAR) with additional UV-A- or UV-B-enriched light, and to ascertain if such alteration is due to known UV-induced stress factors such as reduction of photosynthetic capacity (Jordan, Strid & Wargent, 2016), ROS formation (Hideg et al., 2013), DNA damage (Kalbin et al., 2001) or changes in hormonal status (Hideg & Strid, 2017). The study was carried out in cucumber (*Cucumis sativus* L.) a model species representing broad leaved, high biomass plants of considerable economic importance and which develops considerable phenotypic changes dependent on different wavelengths of UV (Ballaré, Barnes & Kendrick, 1991).

## 2. Materials and Methods

### 2.1. Plant material, growth conditions, and treatment conditions

Cucumber seeds (*Cucumis sativus* L. cv. ‘Hi Jack’) were sown one seed per 0.25 L pot in 14-7-15 NPK fortified peat (SE Horto AB, Hammenhög, Sweden), as described previously (Qian et al., 2019; 2020). Seedlings were grown in a greenhouse under natural daylight from the roof which was supplemented with 150-200 μmol m^−2^ s^−1^ PAR as measured 20 cm above the table using Vialox NAV-T Super 4Y high-pressure sodium lamps (Osram, Johanneshov, Sweden) for 16 h per day centered around solar noon, and only turned off when the natural irradiance reached 900 μmol m^−2^ s^−1^. The day/night temperature was 25/20 °C and the relative humidity was set to 80 %. Watering was done by adding water to the tray underneath the pots when the tray itself was completely dry. As soon as the cucumber seedlings had fully developed cotyledons, watering was commenced using a full nutrient solution (Svegro AB, Ekerö, Sweden).

Fourteen days after sowing, when the first true leaf of the cucumber seedlings was approximately 5 cm in diameter (about one third of the diameter of a fully developed first true leaf), UV exposure commenced. The plants were then given either supplementary UV-A-enriched or UV-B-enriched irradiation for 4 h per day (centered around solar noon) in addition to the PAR described above. Controls were simultaneously exposed to PAR only (see below) in the same chamber as the corresponding UV-treated plants. The UV-A and UV-B exposures were carried out in separate greenhouse chambers and the treatments alternated between the chambers when repeating the experiment (cf. Qian et al., 2019; 2020, for details).

Open top, front and backside boxes (OTFB boxes), covered with Perspex on the left and right sides, were used for the different UV exposures. Each greenhouse compartment was equipped with up to six boxes, three being used for the UV treatments and three for the corresponding controls. Each OFTB box contained up to 48 plants per replicate. For the UV-A-enriched experiments, fluorescent UVA-340 tubes (Q-Lab, Cleveland, Ohio) were used for exposure, whereas for the UV-B-enriched experiments fluorescent Philips TL40/12 UV tubes (Eindhoven, The Netherlands) were employed. For the control OFTB boxes, UV-blocking Perspex was used to cover the top and all sides. For the UV-B-enriched experiment, 0.13mm cellulose acetate (Nordbergs Tekniska AB, Vallentuna, Sweden) covered the top, front and backside of the OTFB boxes with the purpose to remove any UV-C radiation emitted by the Philips TL40/12 tubes. For the UV-A-enriched experiment, the OFTB boxes were similar to the boxes used in the control experiment but without any filtering material on top.

The spectral distribution of the light environments in the different treatments was measured using an OL756 double monochromator spectroradiometer (Optronic Laboratories, Orlando, FL, USA) 20 cm above the table. The details of doses were as described by Qian et al. (2019). Briefly, however, UV-A-enriched radiation contained 3.6 W UV-A m^−2^ and a 45.5 mW m^−2^ plant-weighted UV-B (calculated according to Yu & Björn, 1997), giving a total of plant-weighted UV-B of 0.6 kJ m^−2^ day^−1^ during the daily four-hour exposure. The UV-B-enriched irradiation had 83.4 mW m^−2^ plant-weighted UV-B totaling 1.23 kJ m^−2^ day^−1^ plant-weighted UV-B. This exposure also contained 0.34 W UV-A m^−2^. The daily irradiation outside in Lund, Sweden, under clear skies on a summer’s day is approximately 4.8 kJ m^−2^ day^−1^ of plant-weighted UV-B (Yu & Björn, 1997).

### 2.2. Morphological measurements

Between day 0 and day 14 morphological parameters were measured. A ruler was used to measure the lengths of stems and petioles. The dry matter (DM) of shoots (separated into stems, petioles, and leaves) and roots was measured using a digital balance (accuracy 0.001 g) following oven drying at 70 °C for 20 h. The leaf mass fraction (LMF) was calculated as LMF = leaf DM/total shoot DM. The leaf area (LA) was determined from digitized photographs using ImageJ (https://imagej.nih.gov/ij/). As a measure of how much leaf area a plant builds with a given amount of leaf biomass, the specific leaf area (SLA) for true leaves 2, 3, and 4 from the base of the plans, as well as for all leaves combined, was calculated as SLA=LA/leaf DM. For each experiment, six plants per treatment were measured, two from each of the three replicated treatment OTFB boxes and their corresponding controls. In total three independent experiments were performed.

### 2.3. Chlorophyll fluorescence

Chlorophyll *a* fluorescence was measured with a MINI-PAM (Walz, Effeltrich, Germany) on the 1^st^ true leaf from the bottom of the stem on day 1, 4, 8 and 14 of UV exposure, as well as on the youngest well-developed leaf (the top leaf which was fully developed, i.e. the diameter reached approximately 15 cm) on day 15 of UV exposure. The attached leaves were fixed in a leaf clip holder (2030-B, Walz, Effeltrich, Germany) fitted with a halogen lamp (2050-HB, Walz, Effeltrich, Germany) and a heat absorbing glass filter (Calflex, Optic Balzers, Liechtenstein, Germany). The leaf was dark adapted 30 min by aluminium foil and the middle portion of the leaf was mounted in the leaf clip in a dark room. F_o_ and F_m_ were measured and the maximum photochemical efficiency of photosystem II (PSII) was calculated as F_v_/F_m_ = (F_m_-F_o_)/F_m_. Subsequently, the leaf was exposed to actinic PAR of 302 or 1860 μmol m^−2^ s^−1^ for 10 mins to achieve steady-state F_s_ and F_m_’, measured by 0.6 s saturating pulses. The operation efficiency of PSII (F_q_’/F_m_’), where F_q_’ = F_m_’-F_s_, the fraction of open PSII expressed as q_L_, and non-photochemical quenching (NPQ) were calculated as reviewed by Murchie and Lawson (2013). For each experiment, three plants per treatment were measured (one from each of the three replicated treatment OTFB boxes with accompanying control boxes) and in total three independent experiments were performed.

### 2.4. Biochemical analysis

Upper surface chlorophyll content in the 1^st^ true leaf (on day 1, 4, 8, and 14 of UV exposure), and the youngest well-developed leaf on day 15 of UV exposure was measured using a DUALEX^®^ SCIENTIFIC (Force-A, Orsay, France), following chlorophyll fluorescence measurements.

Leaf adaxial epidermal flavonol content (LAEFC) in the 2^nd^ true leaf (on day 0, 1, 3, 5, 10, and 14 of UV exposure) was measured using a DUALEX (Force-A, Orsay, France). Additionally, total UV-absorbing pigments (TUAP; mostly flavonoids) were extracted from the 2^nd^ true leaf (on day 0, 1, 3, 5, 10, and 14 of UV exposure) for quantification. Leaves were snap frozen, then stored at −80°C until used. They were then ground in liquid nitrogen using a mortar and pestle and 0.1 g leaf material was placed into micro-tubes with 1 ml acidified methanol (1% HCl, 20% H_2_O, 79% CH_3_OH) before incubation in the dark at 4°C for four days. Absorbance was recorded at 330nm using a spectrophotometer (Shimadzu UV/VIS 1800). Absorbance was normalized per leaf fresh weight. For each experiment, three plants per treatment were measured (one from each of the three replicated treatment OTFB boxes with accompanying control boxes) and in total three independent experiments were performed.

With the same sampling as for flavonoid analysis, total antioxidant capacity (TAC) was analyzed using a commercially available kit (Total Antioxidant Capacity Assay kit, Sigma-MAK187). Ground leaf tissue (0.1 g; see above) from the 2^nd^ true leaf was extracted in 1 ml of ice cold 1 X Phosphate Buffered Saline (PBS) and following centrifugation, the supernatant was diluted 1:100 to bring values within range of kit standards. Samples were assayed according to the manufacturer’s protocol, by comparing the absorbances of diluted extracts at 570 nm with Trolox standards and values normalized to tissue fresh weight.

### 2.5. DNA damage detection

With replications as for flavonoid analysis, cyclobutane pyrimidine dimers were quantified using an immunoassay following the protocol from van de Poll, Eggert, Buma & Breeman (2001). First, DNA was extracted from ground leaf tissue (see above) from the 2^nd^ true leaf from the base of the plant on day 1, 3, 5, 10, and 14 using the E.Z.N.A.^®^ Plant DNA Kit (Omega Bio-Tek, Georgia, USA), and dissolved in 100 μl TE buffer (pH 8.0). The DNA concentration was determined fluorometrically in a microplate reader (GENius, Tecan, Salzburg, Austria) using the Quantifluor dye (Promega Madison, USA). Of each sample 50 ng DNA was used for the southern blot. CPDs were subsequently labelled on the membrane by using a primary antibody against CPDs produced in mouse (H3 clone 4F6, Sigma Aldrich, St. Louis, USA). Detection was conducted by peroxidase coupled to the secondary antibody (Anti-Mouse IgG (whole molecule)-Peroxidase, Sigma Aldrich) using an enhanced chemiluminescent substrate (Pierce ECL, Thermo Fisher Scientific, Waltham, USA). On each blot a CPD calibration standard was included to allow absolute quantification of CPDs Mb^−1^ (Pescheck et al. 2014).

### 2.6. Plant hormone analysis

The 2^nd^ true leaves from the base of the plant were harvested on day 3 and day 5. Leaves from three different plants within one experiment were pooled to obtain approx. 300 mg material. Leaf tissues were snap frozen in liquid nitrogen, and kept at −80°C. All samples were then ground with a mortar and pestle in liquid nitrogen, and again kept at −80°C until used. Totally, three independent experiments generated three replicates. Each pooled sample was subdivided in separate 100 mg fractions for the extraction of the different hormone groups.

#### Auxin and ABA analysis

Samples were extracted in 500 μL of 80% methanol. [C^13^]-IAA (100 pmol, (phenyl-^13^C_6_)-indole-3-acetic acid, 99%, Cambridge Isotopes, Tewksbury, MA, USA) and D6-ABA (150 pmol, [^2^H_6_](+)-*cis,trans*-abscisic acid, [(S)-5-[^2^H_6_](1-hydroxy-2,6,6-trimethyl-4-oxocyclohex-2-en-1-yl)-3-methyl-(2Z,4E)-pentadienoic acid], Olchemim, Olomouc, Czech Republic) were added as internal tracers. After overnight extraction, samples were centrifuged (20 min, 15,000g, 4°C, in an Eppendorf 5810R centrifuge, Eppendorf, Hamburg, Germany) and the supernatants were aliquoted in two equal parts. One aliquot was acidified using 5.0 mL of 6.0% formic acid and loaded on a reversed-phase (RP)-C18 cartridge (500 mg, BondElut Varian, Middelburg, The Netherlands). The compounds of interest (IAA, ABA, and the oxidation products IAA-OX, IAA-OH, indole-butyric acid (IBA)-OX, and IBA-OH) were eluted with 5.0 mL of diethyl ether and dried under a nitrogen stream (TurboVap LV Evaporator, Zymark, New Boston, MA, USA). The remaining aliquot was hydrolyzed in 7.0 M NaOH for 3 h at 100°C under a water-saturated nitrogen atmosphere. After hydrolysis, the samples were acidified using 2.0 M HCl, desalted on an RP-C18 cartridge (500 mg), and eluted with diethyl ether. All samples were methylated using ethereal diazomethane to improve analysis sensitivity. Samples were analysed using an Acquity UPLC system linked to a TQD triple quadrupole detector (Waters, Milford, MA, USA) equipped with an electrospray interface in positive mode. Samples (6.0 μL) were injected on an Acquity UPLC BEH C18 RP column (1.7 μm, 2.1 × 50 mm, Waters) using a column temperature of 30°C and eluted at 0.3 mL min^−1^ with the following gradient of 0.01 M ammonium acetate (solvent A) and methanol (solvent B): 0−2 min isocratic 90/10 A/B; 2−4 min linear gradient to 10/90 A/B. Quantitative analysis was obtained by multiple reactant monitoring of selected transitions based on the MH^+^ ion (dwell time 0.02 s) and the most appropriate compound-specific product ions in combination with the compound-specific cone and collision settings. All data were processed using Masslynx/Quanlynx software V4.1 (Waters). Data are expressed in picomoles per gram fresh weight (pmol g^−1^ FW).

#### Gibberellin analysis

Samples were extracted overnight in 500μL acidified methanol pH 4.0 (80/20, methanol/5.0 mM formic acid-containing butylated hydroxytoluene (3−5 crystals)). As internal tracers, D_2_-GA1 (C_19_H^222^H_2_O_6_), D_2_-GA4 (C_19_H_22_^2^H_2_O_5_), D_2_-GA8 (C_19_H_22_^2^H_2_O_7_), D_2_-GA9 (C_19_H_22_^2^H_2_O_4_), D_2_-GA15 (C_20_H_24_^2^H_2_O_4_), D_2_-GA19 (C_20_H_24_^2^H_2_O_6_), D_2_-GA20 (C_19_H_22_^2^H_2_O_5_), and D_2_-GA29 (C_19_H_22_^2^H_2_O_6_) (20 pmol each, Olchemim) were added. After purification on an RP-C18 cartridge (500 mg) as described above for auxins, samples were derivatized with N-(3-dimethylaminopropyl)-N′-ethylcarbodiimide hydrochloride (Sigma-Aldrich, 1.0 mg per sample, pH 4.0, 60 min, 37 °C under continuous shaking in an Eppendorf thermomixer) and analysed using a UPLC-MS/MS equipped with an electrospray interface in positive mode (ACQUITY, TQD, Waters). Samples (6.0 μL, partial loop mode using a 10 μL sample loop) were injected on an ACQUITY BEH C18 column (2.1 × 50 mm; 1.7 mm, Waters) using a column temperature of 30 °C and eluted at 450 μL min^−1^ with the following gradient of 0.1% formic acid in water (solvent A) and 0.1% formic acid in acetonitrile (solvent B): 0−0.8 min isocratic 92/8 A/B; 0.8−5 min linear gradient to 60/40 A/B; 5−5.5 min linear gradient to 10/90 A/B. Quantitative analysis was performed by multiple reactant monitoring of selected transitions based on the MH^+^ ion (dwell time 0.02 s) and the most appropriate compound-specific product ions in combination with the compound-specific cone and collision settings. Transitions are grouped in specific time windows according to the compound-specific retention time in order to keep the dwell time at 0.02s. All data were processed using Masslynx/Quanlynx software V4.1 (Waters). Data are expressed in picomoles per gram fresh weight (pmol g^−1^ FW).

### 2.7. Statistical analysis

Statistical analysis was performed using either SPSS 19.0 (IBM, Armonk, NY), STATA 14.0 or Wizard for Macintosh (App Store, Apple Inc., Cupertino, CA). Morphological parameters (Figs. 1-5) were analyzed using error propagation where the standard deviation of the ratios were approximated using Taylor linearization (Taylor, 1997) as further described in Qian et al. (2020), and tests of differences between means due to treatment (UV-A, UV-B or control) were performed using analysis of variance (ANOVA). Paired T-tests were used to test for changes in length of 1^st^-7^th^ petioles length and parameters related to the 2^nd^-4^th^ true leaves (Figs. 2-4; Tables 1 and 2). For data generated at different time periods of UV exposure, including analysis of chlorophyll fluorescence, chlorophyll content of the 1^st^ true leaf, flavonoid content, TAC, and DNA dimer, two-way ANOVA was performed to test whether each variable was significantly affected by treatment or time of UV exposure. For data including chlorophyll fluorescence and chlorophyll content of the youngest well-developed leaf, and plant hormone concentration, T-tests were performed to analyze if the differences between UV-exposed and control samples were significant or not. Statistical analysis was performed using either SPSS 19.0 (IBM, Armonk, NY) or Wizard for Macintosh (App Store, Apple Inc., Cupertino, CA)

**Figure 1.**
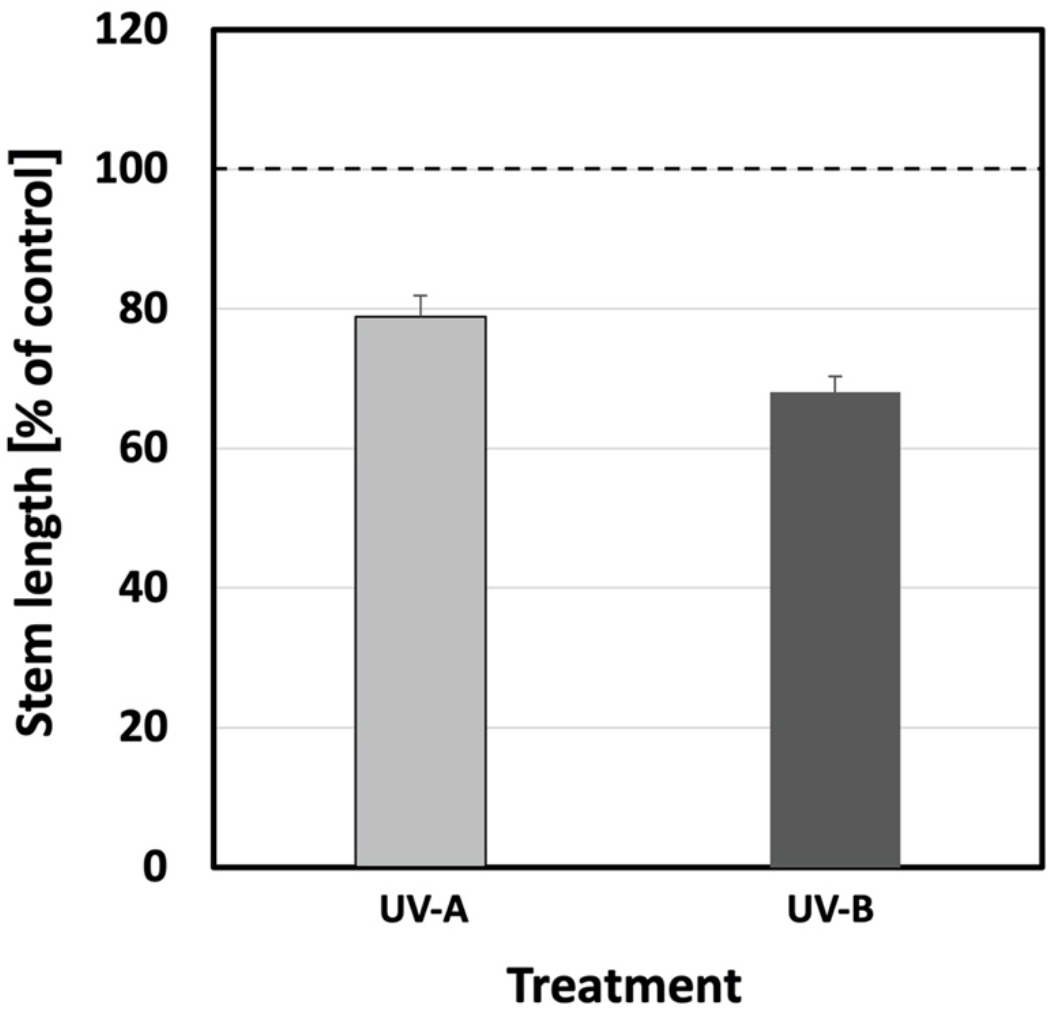
The relative change in stem length of cucumber plants grown under UV-A-enriched (light grey) or UV-B-enriched (dark grey) light, respectively, compared with the corresponding controls. The data represent mean values with n=18 for all treatments and controls ± estimated 95 % confidence interval (whiskers) of the ratio obtained from the approximated standard deviation which in turn was obtained by Taylor linearization (Taylor, 1997). The pairwise comparisons UV-A:control, UV-B:control, and UV-A:UV-B, were all significant (p<0.05; see Table 1).

**Figure 2.**
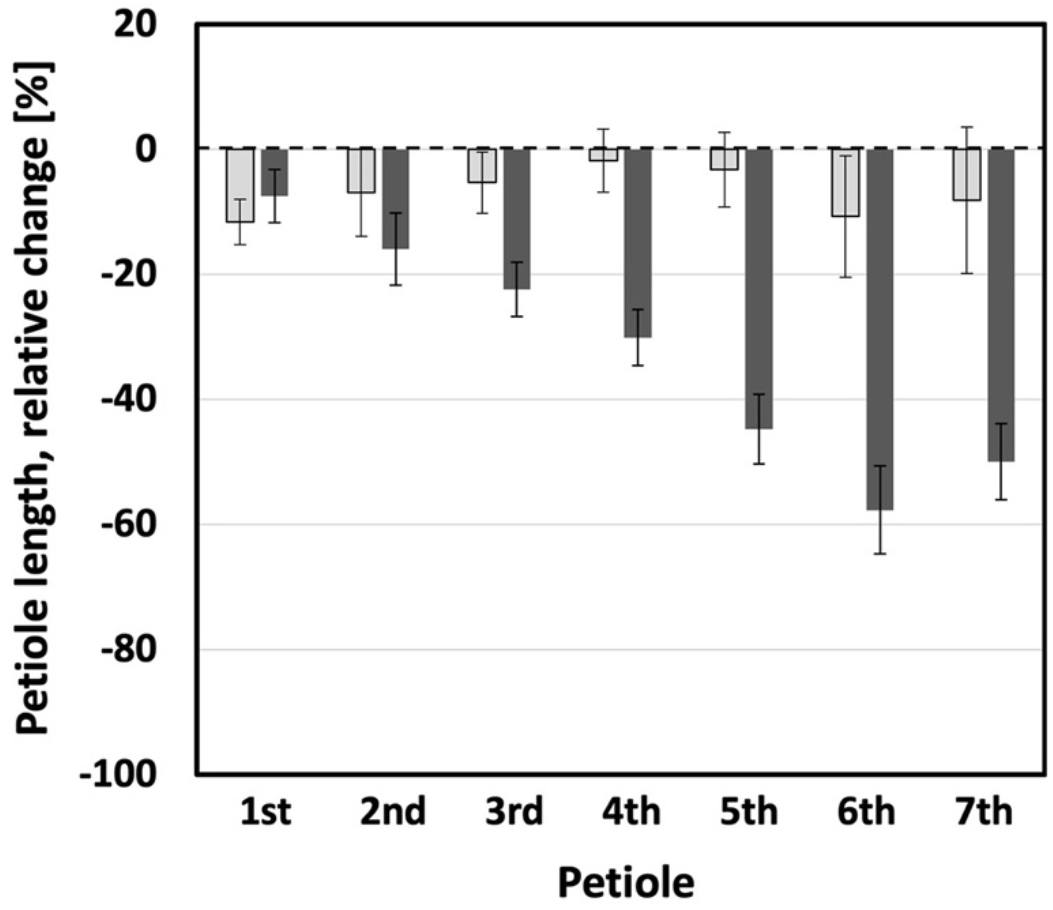
The relative decrease of the lengths of the 1^st^ to 7^th^ petioles of cucumber plants when grown 14 days under UV-A-enriched (light grey) or UV-B-enriched light (dark grey), respectively, compared with the corresponding controls. The data represent mean values with n=18 for all treatments and n=36 for controls ± 95% confidence interval obtained using the approximated standard deviation which was obtained by Taylor linearization (Taylor, 1997). The significant differences (p<0.05) of the pairwise comparisons UV-A:control, UV-B:control, and UV-A:UV-B, are shown in Table 1.

**Table 1.** Significant differences between means of measured variables with regards to treatment and where ‘x’ denotes p<0.05. T-tests were used for total plant or leaf parameters, whereas paired t-tests were used for petiole and true leaf parameters comparing developmental effects on same plant individuals. Petioles and leaves are numbered in order of appearance, with higher numbers for younger structures.

| Plant part | Parameter | UV-A : control | UV-B : control | UV-A : UV-B | Fig. no. |
| --- | --- | --- | --- | --- | --- |
| Stem | Length | x | x | x | 1 |
| 1<sup>st</sup> petiole | Length | x | x | x | 2 |
| 2<sup>nd</sup> petiole | Length | x | x |  | 2 |
| 3<sup>rd</sup> petiole | Length | x | x | x | 2 |
| 4<sup>th</sup> petiole | Length |  | x | x | 2 |
| 5<sup>th</sup> petiole | Length |  | x | x | 2 |
| 6<sup>th</sup> petiole | Length |  | x | x | 2 |
| 7<sup>th</sup> petiole | Length |  | x | x | 2 |
| 2<sup>nd</sup> true leaf | Area | x | x |  | 3A |
| 3<sup>rd</sup> true leaf | Area |  | x | x | 3A |
| 4<sup>th</sup> true leaf | Area |  | x | x | 3A |
| Total leaf | Area | x | x | x | 3B |
| 2<sup>nd</sup> true leaf | Dry mass |  | x | x | 3C |
| 3<sup>rd</sup> true leaf | Dry mass | x | x | x | 3C |
| 4<sup>th</sup> true leaf | Dry mass | x | x | x | 3C |
| 2<sup>nd</sup> true leaf | SLA | x | x | x | 4A |
| 3<sup>rd</sup> true leaf | SLA | x | x | x | 4A |
| 4<sup>th</sup> true leaf | SLA | x |  | x | 4A |
| Total leaf | SLA | x |  | x | 4B |
| Total plant | Dry mass | x | x | x | 5A |
|  | Shoot/root ratio | x |  |  | 5B |
|  | LMF | x | x | x | 5C |

**Table 2.** Pairwise comparisons for statistical significance (paired t-test) of petiole length and true leaf area in plants exposed to UV-B-enriched light and where * is p<0.05; ** is p<0.01; *** is p<0.005; **** is p<0.001 and n.s. is not statistically significant. Petioles and leaves are numbered in order of appearance, with higher numbers for younger structures.

| Treatment | Parameter | Plant part 1 | Plant part 2 | Level of significance | Fig. no. |
| --- | --- | --- | --- | --- | --- |
| UV-B-enriched | Petiole length | 1<sup>st</sup> petiole | 2<sup>nd</sup> petiole | *** | 2 |
|  |  | 2<sup>nd</sup> petiole | 3<sup>rd</sup> petiole | ** | 2 |
|  |  | 3<sup>rd</sup> petiole | 4<sup>th</sup> petiole | **** | 2 |
|  |  | 4<sup>th</sup> petiole | 5<sup>th</sup> petiole | **** | 2 |
|  |  | 5<sup>th</sup> petiole | 6<sup>th</sup> petiole | **** | 2 |
|  |  | 6<sup>th</sup> petiole | 7<sup>th</sup> petiole | **** | 2 |
| UV-B-enriched | True leaf area | 2<sup>nd</sup> true leaf | 3<sup>rd</sup> true leaf | **** | 3A |
|  |  | 3<sup>rd</sup> true leaf | 4<sup>th</sup> true leaf | **** | 3A |

To describe the relationship between the LAEFC and TUAP measurements, a simple regression model was first fitted. Thereafter additional explanatory variables giving number of days of treatment/leaf age and two dummy variables taking into account treatment (UV-A-enriched, UV-B-enriched, and control) were included, to see if they would contribute in describing the dependent variable LAEFC. The final model included the TUAP variable and the variable Days of Treatment as explanatory variables. The residuals were analyzed and the assumptions behind the model seemed to be fulfilled, and showed no signs of systematic pattern, supporting the choice of model.

The same approach was used when TAC was the dependent variable and as explanatory variables in the full model; TUAP, Days of Treatment, two dummy variables for the treatment (UV-A, UV-B, and control). The final model included the same explanatory variables as the first model, TUAP and Days of Treatment, the other explanatory variables were not significant, thereby not contributing to the explanation of the values of TAC The analysis of the residuals did not show any indication on deviations from the model assumptions.

## 3. Results

A more dwarfed, UV-induced, plant architecture has been observed in many different plant species, following exposure to UV radiation. However, some of the strongest morphological responses have been observed in cucumber (e.g. see Qian *et al*., 2020). In the present study we analyzed the induction of a more dwarfed architecture in *C. sativus* cv. Hi Jack. Two-week old cucumber plants were exposed to UV for 14 days, during which the stem length of control plants increased from, on average, 4.1 cm to 47.9 cm. Plants exposed to UV-A- or UV-B-enriched radiation remained comparatively short and reached just 79% and 68% of control stem length, respectively. (Fig. 1; p<0.05; Table 1).

The UV-mediated decrease in elongation growth was not limited to plant height. A similar impediment of elongation could be observed for petioles. The typical petiole length for a control plant ranged between 1.5 and 11.9 cm for the 7th and the 3rd leaf, respectively.

However, petioles of plants exposed to UV-A- or UV-B-enriched radiation remained considerably shorter (Figure 2). The relative effect of UV-A-enriched radiation was more-or-less constant across the range from older to younger leaves, not exceding more than 17% inhibition. The decrease in petiole length was significant (p<0.05) in UV-A-exposed plants for the 1^st^, 2^nd^ and 3^rd^ petiole only (Table 1). In contrast, the effects of UV-B-enriched radiation were particularly pronounced for the youngest leaves (5^th^, 6^th^, and 7^th^ petiole), with petiole length decreasing by more than 40% compared with control plants. This UV-B-induced decrease was significant for the 1^st^-7^th^ petioles (Table 1). Also, progressive decreases in petiole length under UV-B-enriched radiation were found to be statistically significant when comparing adjoining petioles 1 through 6 (Table 2), i.e. a larger decrease the younger the tissue. However, there was no statistically significant difference in the extent of the decrease when comparing the small, developing petioles 6 and 7.

However, petioles of plants exposed to UV-A- or UV-B-enriched radiation remained considerably shorter (Figure 2). The relative effect of UV-A-enriched radiation was more-or-less constant across the range from older to younger leaves, not exceeding more than 17% inhibition. The decrease in petiole length was significant (p<0.05) in UV-A-exposed plants for the 1^st^, 2^nd^ and 3^rd^ petiole only (Table 1). In contrast, the effects of UV-B-enriched radiation were particularly pronounced for the youngest leaves (5^th^, 6^th^, and 7^th^ petiole), with petiole length decreasing by more than 40% compared with control plants. This UV-B-induced decrease was significant for the 1^st^-7^th^ petioles (Table 1). Also, progressive decreases in petiole length under UV-B-enriched radiation were found to be statistically significant when comparing adjoining petioles 1 through 6 (Table 2), i.e. a larger decrease the younger the tissue. However, there was no statistically significant difference in the extent of the decrease when comparing the small, developing petioles 6 and 7.

Leaf area was also affected by UV. Generally, only small changes in leaf area were obtained following treatment with UV-A-enriched light (Fig. 3A and B). Yet, exposure to UV-B-enriched light led to considerable decreases in leaf area. The UV-B-induced alteration was more pronounced for younger leaves, as is shown in Fig. 3A for true leaves 2, 3 and 4 (15, 24, and 35% smaller leaves, respectively). For the UV-B treatment these changes were all statistically significant (p<0.05), whereas for UV-A only the 10% decrease in leaf area (Fig. 3A) of true leaf no. 1 was statistically significant (Table 1). In addition, the progressive decrease in true leaf area under UV-B-enriched radiation was statistically significant for true leaf 2 compared with leaf 3, and leaf 3 compared with leaf 4 (p<0.001; Table 2), confirming a larger decrease the younger the tissue. At the whole plant level, this resulted in a statistically significant decrease in leaf area by 5% for plants exposed to UV-A-enriched light and by 28% for plants grown under UV-B-enriched light (Fig. 3B). In parallel, UV-B exposure caused a statistically significant decrease in leaf dry weight for true leaves 2, 3 and 4 (Fig. 3C; Table 1), while UV-A exposure caused a small increase.

**Figure 3.**
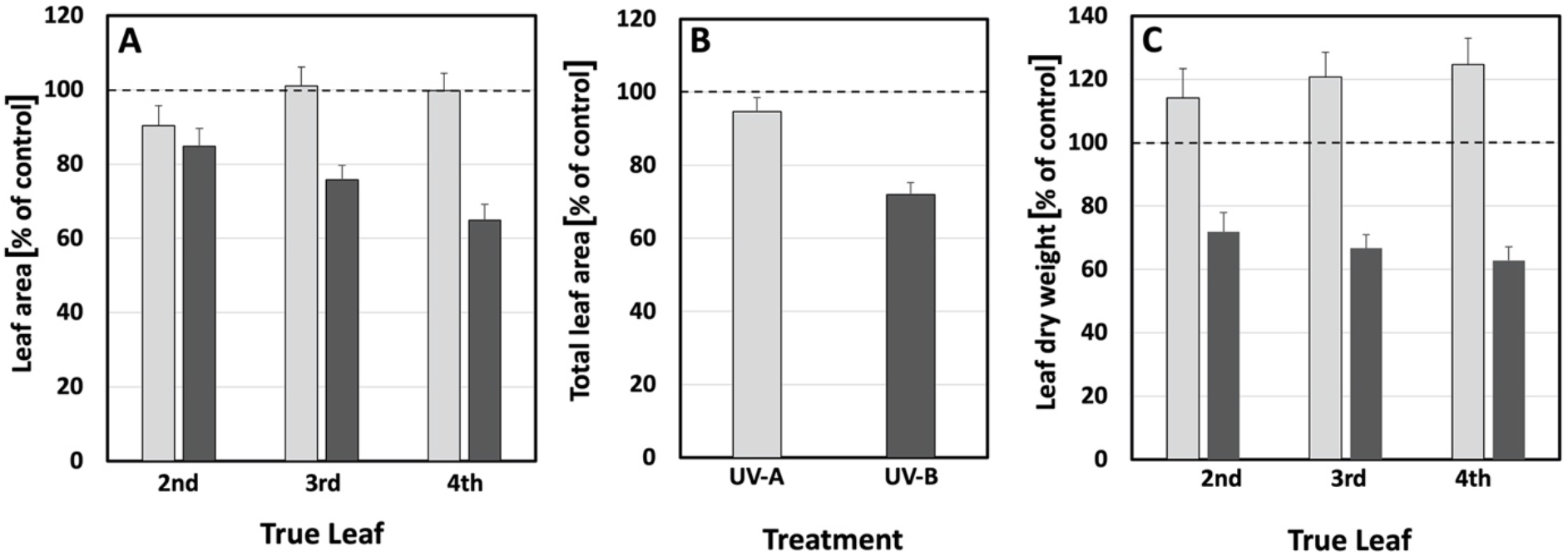
The relative change of (**A**) leaf area of true leaves 2, 3, and 4; (**B**) total leaf area; (**C**) dry weight of true leaves 2, 3, and 4, compared with the corresponding controls when grown under UV-A- or UV-B-enriched light, respectively. The data represent mean values with n=18 for all treatments and n=36 controls ± the estimated 95 % confidence interval (whiskers) of the ratio obtained using the approximated standard deviation which was obtained by Taylor linearization (Taylor, 1997). The significant differences (p<0.05) of the pairwise comparisons UV-A:control, UV-B:control, and UV-A:UV-B,are shown in Table 1.

The observed decrease in leaf area, together with a slightly larger decrease in leaf mass in plants exposed to UV-B-enriched light (Fig. 3A vs 3C), results in a statistically significant increase in specific leaf area (SLA) in the 2^nd^ and 3^rd^ leaves (Fig. 4A and Table 1). The older the leaves, the larger the increase in SLA. A similar trend was also seen in the total plant SLA (Fig. 4B) which, however, was not statistically significant. In contrast, a clear and statistically significant negative effect of UV-A-enriched light on 2^nd^, 3^rd^, and 4^th^ leaf SLA as well as on total plant SLA is discernible, with total SLA decreasing by as much as 18% compared with control leaves (Fig. 4 B and Table 1). There was a statistically significant difference in SLA between plants exposed to UV-A- and UV-B-enriched light in all cases (Fig. 4A and B and Table 1).

**Figure 4.**
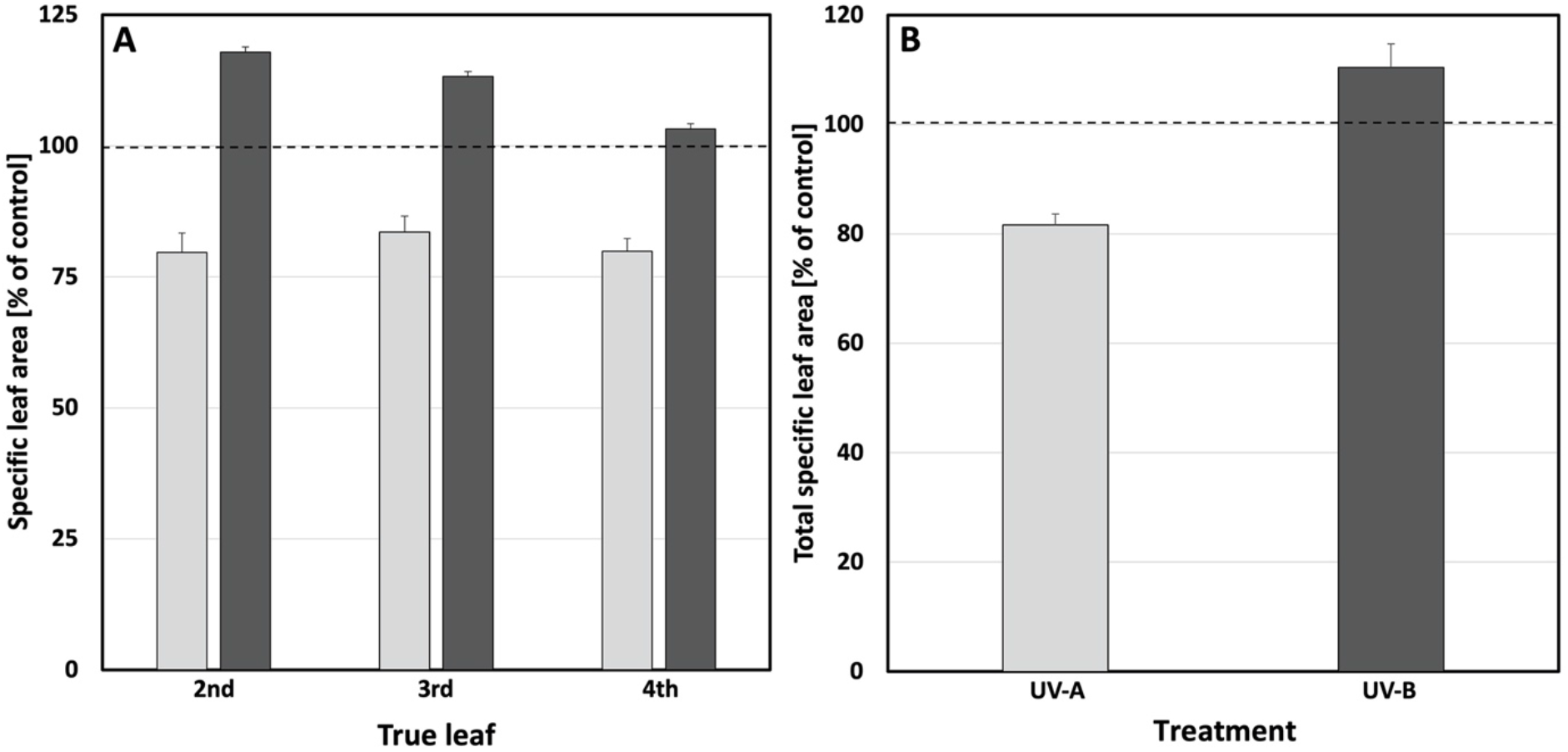
The relative change in (**A**) specific leaf area (SLA [cm^2^ mg^−1^]) of true leaves 2, 3, and 4; (**B**) total SLA compared with the corresponding controls when grown under UV-A-enriched (light grey) or UV-B-enriched (dark grey) light, respectively. The ratios are based on the mean values with n=18 for all treatments and n=36 controls ± estimated 95 % confidence interval (whiskers) obtained using the approximated standard deviation which was obtained by Taylor linearization (Taylor, 1997). The significant differences in specific leaf area (p<0.05) of the pairwise comparisons UV-A:control, UV-B:control, and UV-A:UV-B,are shown in Table 1.

To explore whether the more compact architecture of plants exposed to UV-A- or UV-B-enriched light was related to an overall decrease in growth, both biomass and photosynthetic activity were measured. The dry weight of control plants ranged between 3.8 and 4.6 g (average 4.2 g) after 28 days of growth. Overall, UV-A-enriched radiation significantly stimulated plant biomass production by 14% relative to the control (Fig. 5A and Table 1). In contrast, UV-B-enriched radiation had a clear negative impact on biomass accumulation. The statistically significant decrease in biomass caused by UV-B was 28% (Fig. 5A and Table 1). UV-A also impacted on the shoot-to-root ratio, resulting in a 22% decrease (Fig. 5B and Table 1) due to the relatively high root biomass in plants exposed to UV-A-enriched light. No such effect was seen in plants exposed to UV-B-enriched light. A small but statistically significant increase in the leaf weight fraction, relative to the controls (Fig. 5C and Table 1), was induced by both UV-A and UV-B.

**Figure 5.**
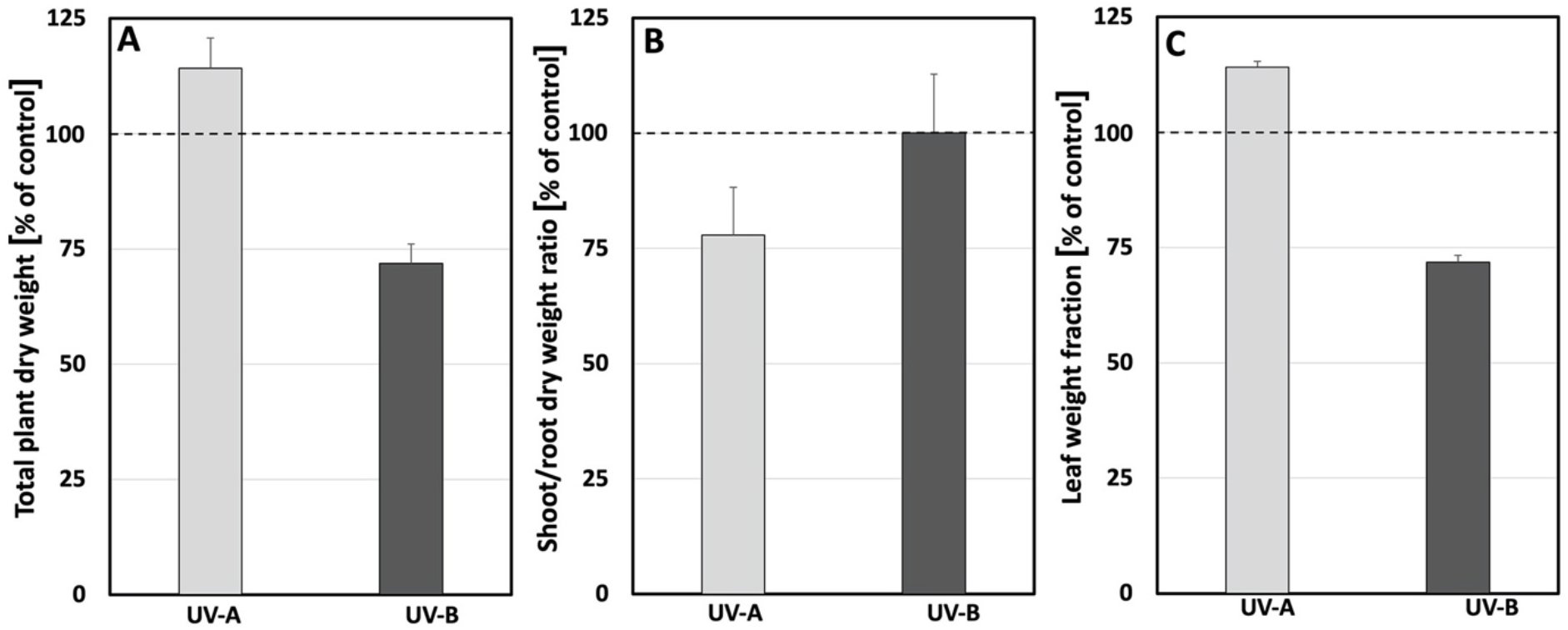
The relative change in (**A**) plant dry matter, (**B**) shoot/root dry matter ratio, and (**C**) leaf weight fraction with the corresponding controls when grown under UV-A-enriched (light grey) or UV-B-enriched (dark grey) light, respectively. The ratios are based on the mean values with n=18 for all treatments and n=36 for controls ± estimated 95 % confidence interval (whiskers) obtained using the approximated standard deviation which was obtained by Taylor linearization (Taylor, 1997). The significant differences (p<0.05) of the pairwise comparisons UV-A:control, UV-B:control, and UV-A:UV-B,are shown in Table 1.

To understand the underlying cause of the observed alterations in plant morphology, it was explored whether UV-exposed plants exhibited disrupted metabolism as a response to stress. Photosynthetic activities were monitored using chlorophyll fluorometry throughout the experiment for all treatments. The initial measurement of F_v_/F_m_ on the four measuring days did not differ between treatment nor day and was 0.792 ± 0.012 (data not shown). Thus, the photosynthetic response did not change over time (data not shown) and just data from day 15, the first day after the UV enrichment, are presented (Fig. 6). The measurements were done on the youngest well-developed leaf and followed by exposure to actinic light at low (302 μmol m^−2^ s^−1^) and high (1860 μmol m^−2^ s^−1^) PAR. The lower PAR corresponded to a level in the range that the plants experienced during most of the day in the greenhouse, while the high level corresponded to light saturation, where potential differences in light acclimation are most clearly shown. Despite the exposed position of the youngest fully developed leaf and exposure to the treatments for the entirety of its development, the operation efficiency of PSII (F_q_’/F_m_’; Fig. 6A) and fraction of open PSII (q_L;_ Fig. 6B) were both unaffected by UV treatments. The only treatment effect was a small but significant increase in heat dissipation through NPQ (Fig. 6C) in cucumbers grown in UV-A-enrichment. This was not a big enough increase to affect F_q_’/F_m_’ and q_L_, indicating that photosynthesis was not affected by the UV-enrichment.

**Figure 6.**
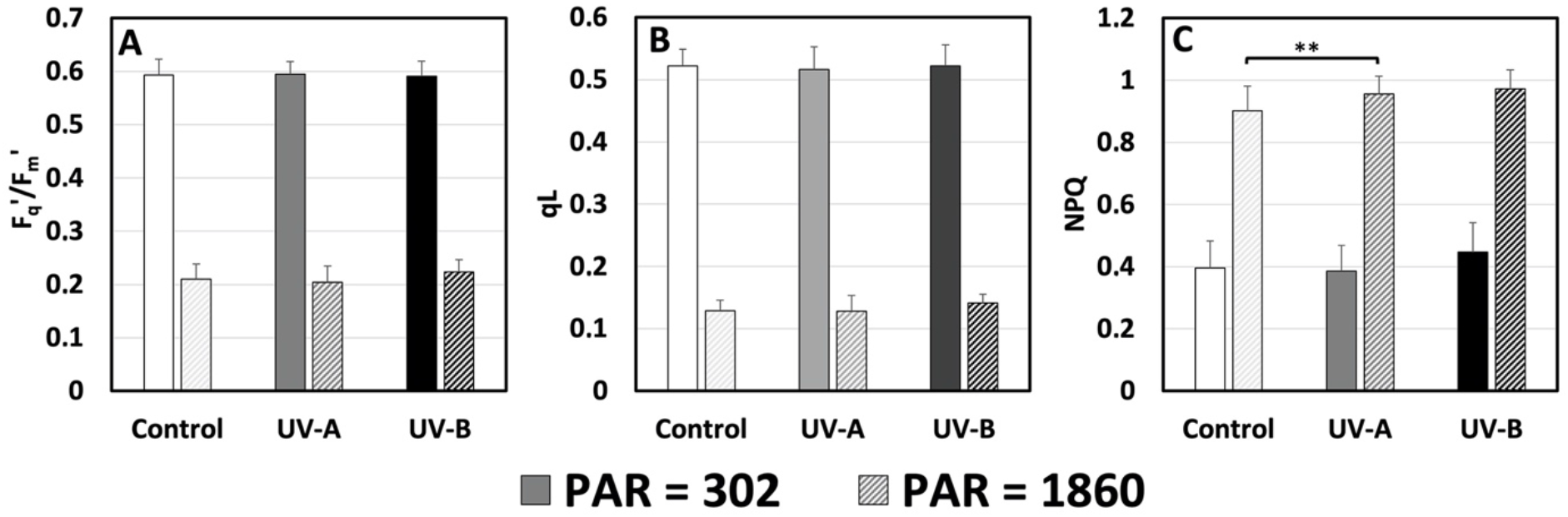
The response of (**A**) the operation efficiency of PSII (F_q_’/F_m_’), (**B**) the fraction of open PSII (q_L_), and (**C**) heat dissipation measured as non-photochemical quenching of fluorescence (NPQ) measured on the youngest well-developed leaf under an actinic PAR of 302 or 1860 μmol m^−2^ s^−1^ day 15 after commencement of UV exposure (last day of UV exposure day 14) to UV-deficient control (white), UV-A-enriched (grey) or UV-B-enriched (black) light. The data represent mean values ± SD with n = 9 for UV treatments and n = 18 for controls. T-test was used for statistical analysis. ** were used to represent the significant difference for P ≤ 0.01.

In parallel to measurements of the photosynthetic activity, chlorophyll content was measured using a Dualex. For the duration of the experiment, the 1^st^ true leaf slowly accumulated more chlorophyll per unit of leaf area (Fig. 7A). Treatment with UV-A- or UV-B-enriched radiation has no impact on this process. Likewise, in the youngest well-developed leaf, measured on day 15 of UV treatment, there was no statistically significant effect of UV-A or UV-B on chlorophyll content (Fig. 7B).

**Figure 7.**
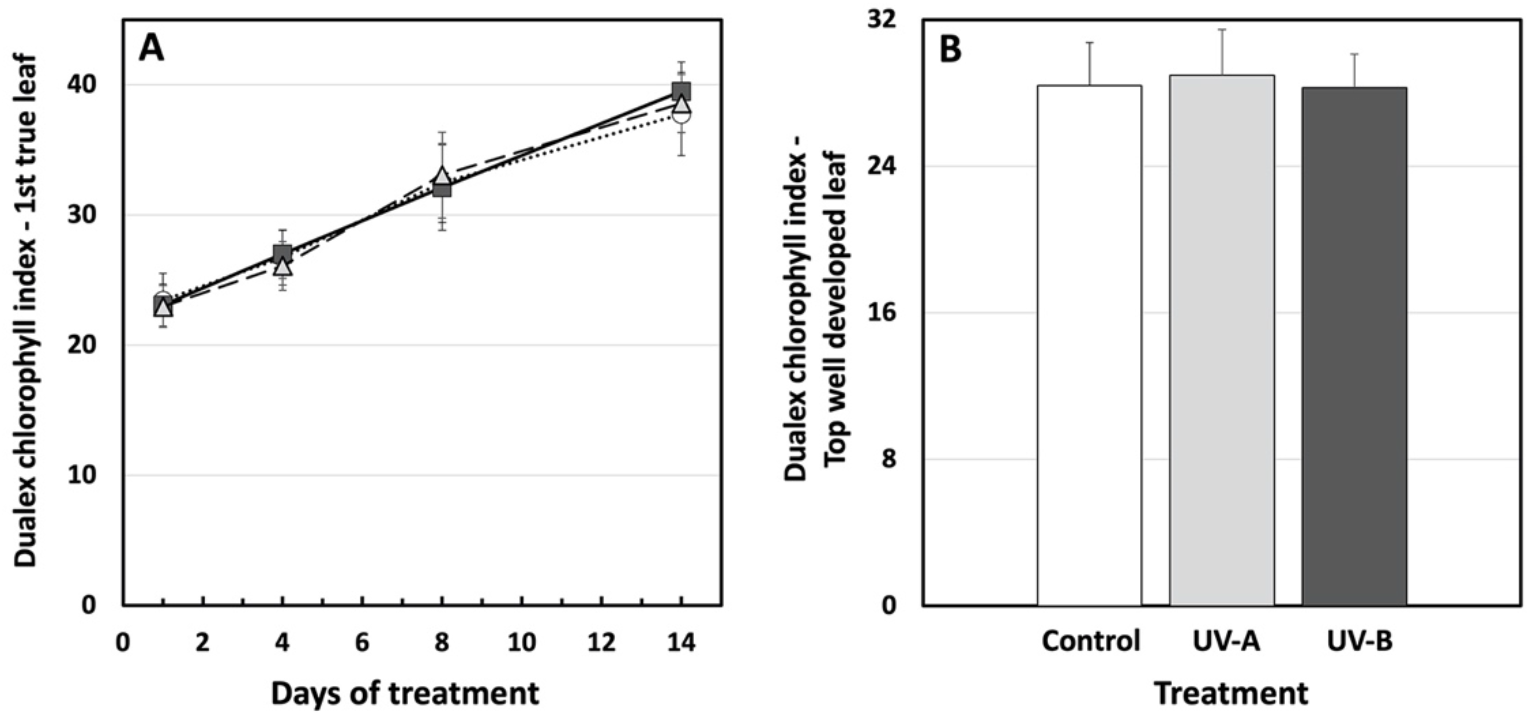
(**A**) The adaxial surface chlorophyll levels expressed as “Dualex chlorophyll index” of the 1^st^ true leaf of UV-deficient controls (open circles), UV-A-enriched light (grey triangles) and UV-B-enriched light (closed squares) treated plants; (B) Adaxial surface chlorophyll levels of the youngest well-developed leaf measured on day 15 after commencement of treatment using UV-A-enriched (light grey) or UV-B-enriched (dark grey) growth light, compared with the corresponding UV-deficient control (white). Sampling was done as in Fig. 1. The data represent mean values ± SD, n=9 for the UV-enriched treatments and n=18 for the control. Two-way ANOVA was used to test the effect of exposure time, and treatment on adaxial surface chlorophyll levels of the 1^st^ true leaf. T-test was used to test the significant difference of upper surface chlorophyll levels in youngest well-developed leaf between UV-treated and control samples.

A key component of plant UV protection is the accumulation of flavonols and related compounds. Here we show a complex induction curve using two independent approaches that both peaked five days after commencement of UV treatment. LAEFC measurements (using the Dualex instrument) revealed that UV-B-enriched, and to a lesser extent UV-A-enriched, radiation induced accumulation of flavonols in the 2^nd^ true leaf (Fig. 8A). The same pattern can be observed in TUAP, reflecting the total leaf content of flavonoids. Here, the absorbance at 330 nm of methanolic extracts of leaf discs from UV-B exposed plants increased substantially. To a lesser extent this was also the case for plants exposed to UV-A-enriched light (Fig. 8B).

**Figure 8.**
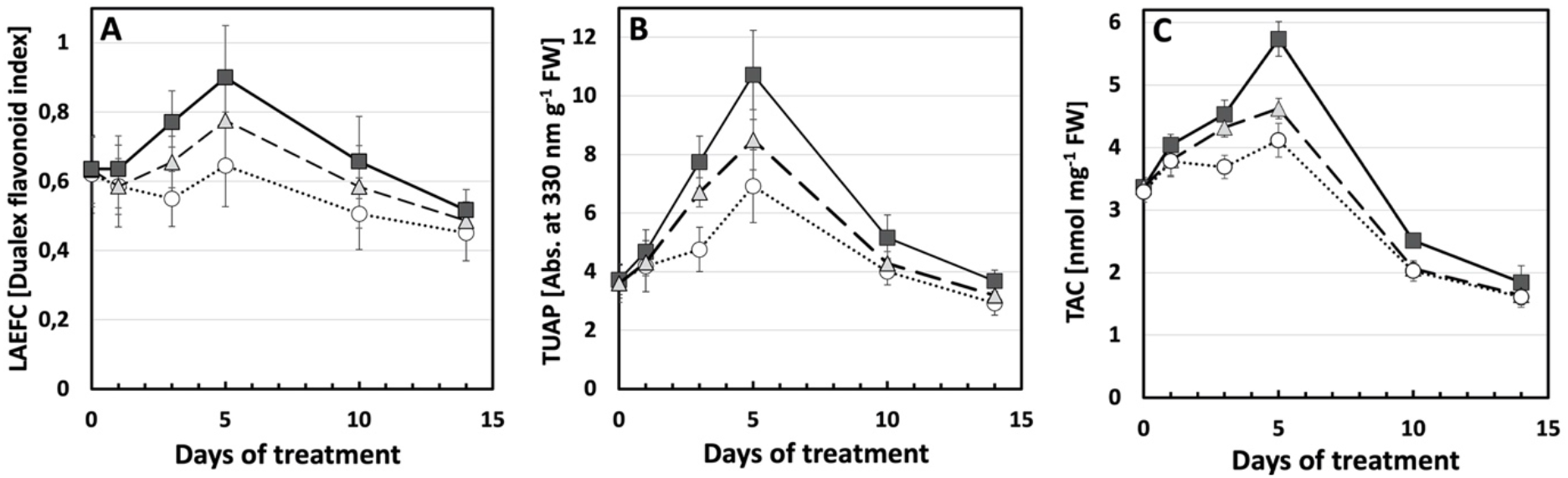
(**A**) Leaf adaxial epidermal flavonol content, LAEFC, measured with Dualex; (**B**) total UV-absorbing pigments, TUAP, measured spectrophotometrically at 330 nm per leaf fresh weight; (**C**) total antioxidant capacity, TAC, measured as nmol Trolox equivalents per mg leaf fresh weight. Measurements were performed on the 2^nd^ true leaf of two-week old cucumber plants grown under UV-A-enriched (grey triangles) or UV-B-enriched light (closed squares), respectively, and compared with the corresponding controls (open circles). Data represent mean values ± SD with n = 9 for the UV-enriched treatments and n = 18 or the control treatment.

To ascertain to what extent the two analytical methods describe the same physiological process within the plant leaf (i.e. that the two different pools of flavonoids measured by these techniques where directly proportional to each other), two models were adapted to describe the relationship between the methods. First, a simple linear relationship between read-outs of the two analytical flavonol assessment methods was assumed (see Supplementary Equation S1 and Supplementary Fig. S1), without taking into account the type of treatment (UV-A- or UV-B-enriched) or leaf age. The R^2^ of this linear fit was 0.76.

In a second model, the dependence between the results of the LAEFC and TUAP methods was assumed to be due also to treatment (UV-A- or UV-B-enriched, or control) and leaf age:

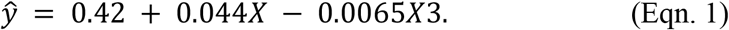

A small influence of leaf age (*X*3) was found. R^2^ of this fit was 0.84, indicating that this model is better in explaining the dependence between LAEFC and TUAP. Also, the residual plots show a random pattern on both sides of 0 (Supplementary Fig. S2A and S2B), thus justifying the model assumptions that leaf age is also a determinant of the relationship between LAEFC and TUAP. Furthermore, and provided that the linear relationship is valid also for x < the observed values, a zero level in the TUAP parameter (i.e. at the intercept of the axis of the LAEFC parameter) corresponded to a Dualex index of 0.42 in Eqn. 1, indicating the presence of flavonols in the epidermal cells when the flavonoid level in the bulk of the leaf is negligible.

Linked to the increase in flavonols, an increase in total antioxidant capacity (TAC) can be seen in leaves exposed to UV-B-enriched light. To a small extent, leaves exposed to UV-A-enriched light also increased their TAC (Fig. 8C). Interestingly, whereas both the LAEFC and TUAP parameters on day 14 returned to a level similar to the one before onset of UV exposure, or slightly below, the TAC parameter decreased to approximately 50% of the initial value, independently of whether the plants had experienced any of the UV exposures or were controls.

To more accurately examine this biphasic nature of the TAC measurements, we studied the dependence between the Trolox assay and the two other methods applied in this study (LAEFC and TUAP). First, for the dependence between the TAC and the LAEFC assays, we assumed two simple linear relationships, without taking into account the type of treatment (UV-A- or UV-B-enriched), but dividing up the samples in those from younger leaves (≤ 5 days of exposure time, i.e. ≤ 19 days after sowing) and those from older leaves (≥ 10 days of exposure time, i.e. ≥ 24 days after sowing). In this case the differences of the linear relationships between the TAC and LAEFC assays became apparent, as is shown in Fig. 9A. The model for the linear dependence with regards to young leaves was found to be:

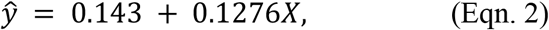

whereas for the linear dependence with regards to older leaves, the equation was:

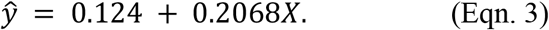

**Fig. 9.**
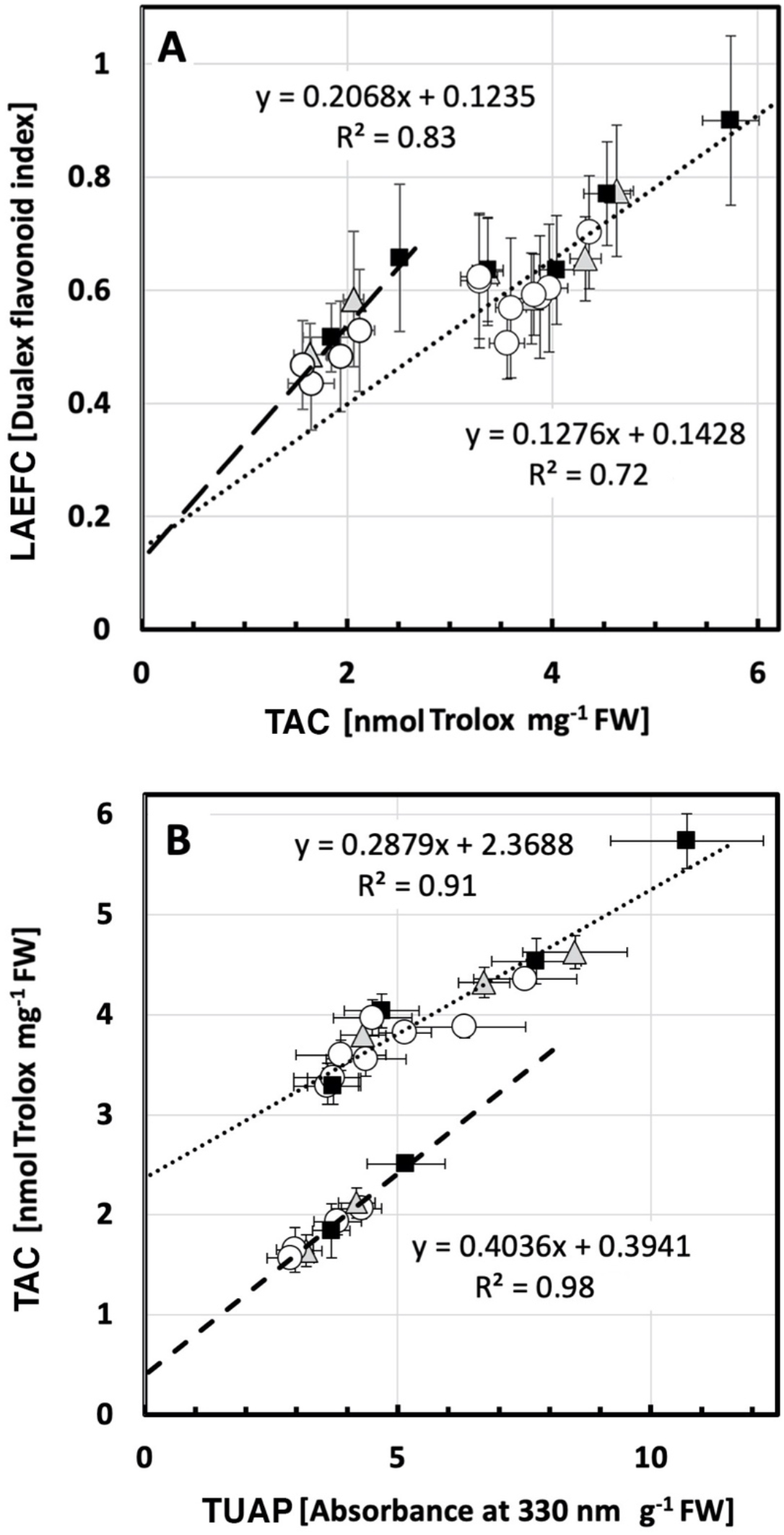
**(A)** The biphasic relationship between TAC and LAEFC (assumed in Eqs. 2 and 3) in UV-deficient controls (open circles), plants grown in either UV-A-enriched light (grey triangles) or UV-B-enriched light (closed squares). The dotted line corresponds to younger leaves (Eqn. 2), whereas the dashed line corresponds to older leaves (Eqn. 3). (**B**) The biphasic relationship between TAC and TUAP (same symbols as in A), assumed in Eqs. 4 and 5. The dotted line corresponds to younger leaves (Eqn. 2), whereas the dashed line corresponds to older leaves (Eqn. 3).

R^2^ for the fits of data points to the two assumed linear relationships was 0.72 and 0.83, respectively. The intercept at zero TAC level was similar for both linearizations with a LAEFC value of approximately 0.13 (Eqs. 2 and 3), indicating a low level of leaf epidermal flavonols that are not active as antioxidants.

We also applied a second model to describe the results of the LAEFC measurements with the TAC method as an explanatory variable and also including explanatory variables treatment (UV-A- or UV-B-enriched) and leaf age (see Supplementary material). However, in this case we did not find any statistically significant proof for the assumption that UV treatment or leaf age influences the correlation (see Supplementary Eqn. S2) and we could not conclude which of the two models that better explained the correlation between data points.

Finally, we scrutinized the linear dependence between the TUAP (Fig. 8B) and the TAC methods (Fig. 8C), using the same two models as applied above. In Fig. 9B, we again assumed two simple linear relationships, without taking into account the type of treatment (UV-A- or UV-B-enriched), but dividing up the samples in those for younger leaves (≤ 5 days of exposure time, i.e. ≤ 19 days after sowing) and those from older leaves (≥ 10 days of exposure time, i.e. ≥ 24 days after sowing). Two linear relationships between the TAC and TUAP data were estimated and the equation with regards to younger tissue was found to be:

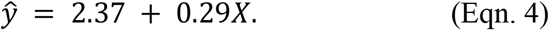

In this case the equation explained variation in data with a factor R^2^ of 0.91. For older tissue the equation became:

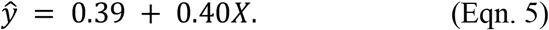

R^2^ was even higher (0.98) for the older leaves and the difference between the intercepts at 0 absorbance for TUAP differed approximately 6-fold (0.39 to 2.37). A second model, where involvement of both treatment effects (UV-A- or UV-B-enriched) and leaf age was considered, was also tested. Now, a clear effect of tissue age was seen on the dependence between TUAP and TAC (Supplementary Eqn. S3). Thus, leaf age has to be considered when comparing results of assays for leaf total flavonoids and anti-oxidative capacity.

To further understand the link between UV exposure and induced changes in plant morphology, accumulation of CPD dimers was measured in leaves of plants exposed to UV-A- or UV-B-enriched light and in control plants to see whether induced CPDs were associated with the smaller cucumber phenotype. Overall, the number of CPDs was low, and, with the exception of day 14, variability was limited. There were no statistically significant effects of UV-A or UV-B on CPD accumulation (Fig. 10).

**Figure 10.**
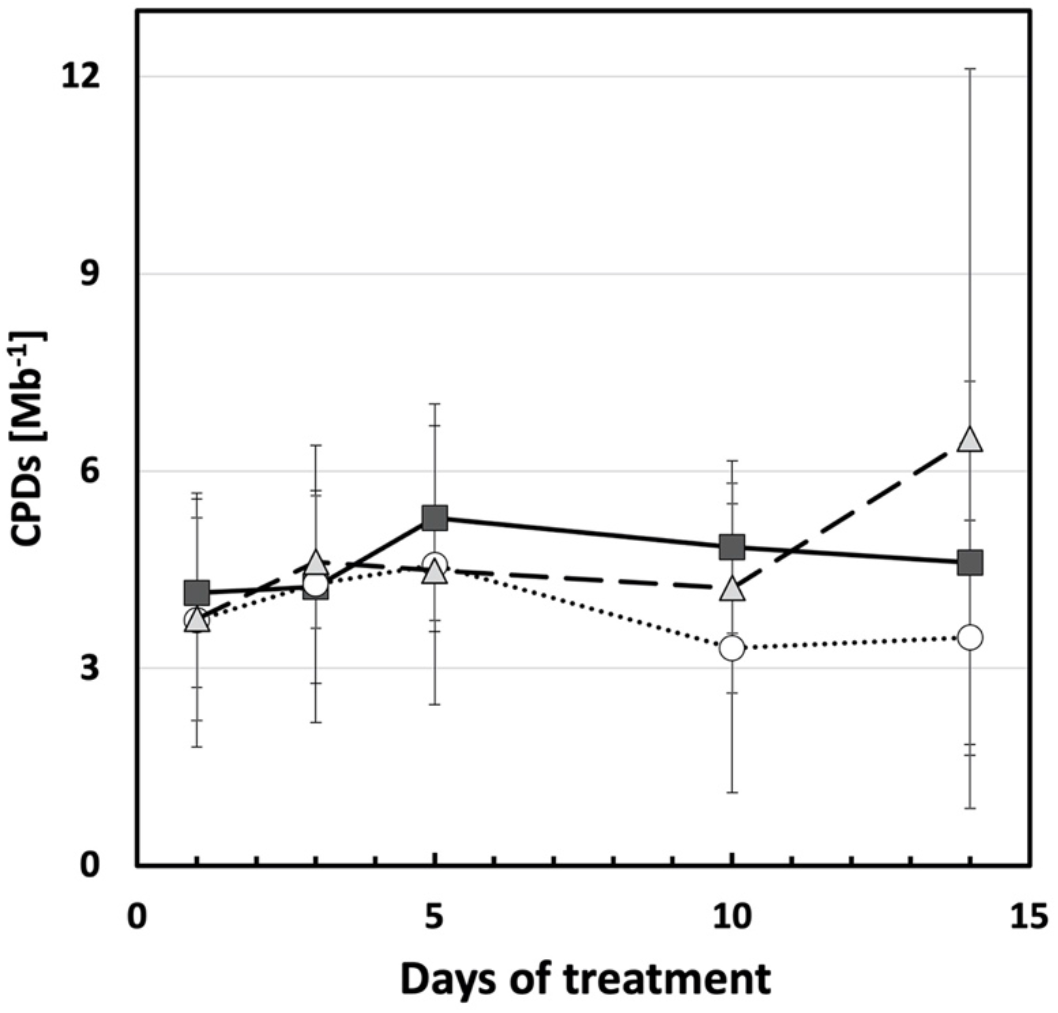
Cyclobutane pyrimidine dimers per mega base (CPDs Mb^−1^) of the 2^nd^ true leaf during the UV treatments compared with the corresponding controls (open circles) when grown under UV-A-enriched (grey triangles) or UV-B-enriched (closed squares) light, respectively. The data represent mean values ± SD with n = 9 for the UV-enriched treatments and n = 18 or the control treatment. Two-way ANOVA was used to test the effect of exposure time and treatment.

Finally, to explore whether changes in key plant hormones are associated with observed changes in plant morphology, leaf concentrations of abscisic acid (ABA), gibberellins (GA) and the auxins indole acetic acid (IAA) and indole butyric acid (IBA), were quantified. Statistically significant decreases in ABA, IAA, and IBA were associated with leaf development (Table 3). No statistically significant effects of UV treatment were observed, although small decreases in the gibberellins GA1, GA44, GA6 and GA15 were noted in plants exposed to either UV-A or UV-B enriched radiation (Table 3).

**Table 3.**
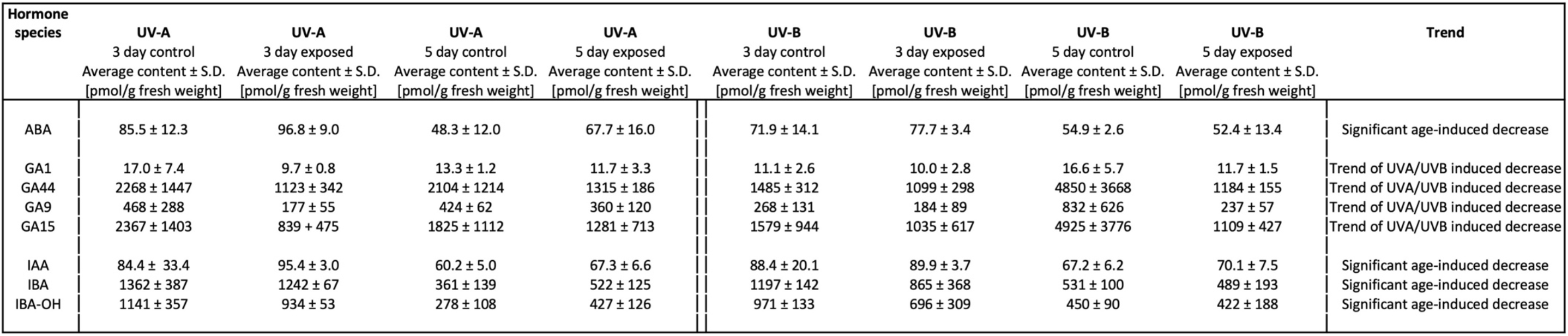
Main plant hormones, including abscisic acid (ABA), gibberellins (GA), and the auxins indole acetic acid (IAA) and indole butyric acid (IBA)), present in the 2^nd^ true leaf on day 3, and 5 of UV treatments. Plant tissue from one plant in each of three boxes per treatment was pooled together with three separate experiments giving n = 3 for both the UV-enriched treatments and the control treatment. The data represent mean values ± SD. For data of each plant hormone species, T-test was used to test the significant difference between each two treatments. The levels of the following hormone species were under the detection limit: GA3, GA4, GA5, GA8, GA12, GA19, GA20, IAA-OX, IAA-OH, IBA-OX, IBA-OH-C, IBA-OX-C. The levels of the following hormone species did not show any clear trend: GA7, IAA-C, IBA-C.

## 4. Discussion

### 4.1. Cucumber displays a strong morphological response when exposed to supplemental UV

Here we explored the regulation of plant morphology by white light enriched with either UV-A or UV-B. The data show that enrichment of the spectrum with either UV wavelength results in a stocky cucumber phenotype. These data are in agreement with previous studies that show that cucumber is particularly responsive to UV-exposure (Krizek et al., 1978; Murali & Teramura, 1986; Ballaré et al., 1991; Adamse & Britz, 1992; Adamse, Britz & Caldwell, 1994; Krizek, Mirecki & Kramer, 1994; Takeuchi, Kubo, Kasahara & Sasaki, 1996; Krizek, Mirecki & Britz, 1997; Fukuda, Satoh, Kasahara, Matsuyama & Takeuchi, 2008; Shinkle, Edwards, Koenig, Shaltz & Barnes, 2010; Yamasaki, Shimada, Kuwano, Kawano & Noguchi, 2010; Yamasaki, Shigeto, Ashihara & Noguchi 2014; Qian et al., 2019, 2020). Therefore, we consider cucumber a promising model species for the study of the stocky UV phenotype, a role that will be facilitated by the large leaf surface area which enables measurement of potential hormone gradients within organs. The data presented in this study show that especially elongation is affected by UV-exposure, with noted decreases in stem and petiole length, as well as in leaf area. These effects were larger in plants that had been grown in the UV-B-enriched light environment than in the UV-A-enriched dito (Figs. 1, 2, 3A and 3B). Interestingly, whereas the petiole lengths and leaf area in plants treated with UV-A-enriched light were approximately the same independently of tissue age, there was a progressively larger decrease in these parameters the younger the tissue in the presence of UV-B radiation (Figs. 2 and 3A; Table 2). Finally, in plants grown under the UV-A-enriched light regimen, a significant reallocation of photosynthate from shoot to root by more than 20% was seen (Fig. 5B). The differences between the effects of UV-A- and UV-B-enriched light on cucumber morphology indicate that there may be different developmental regulatory mechanisms involved in the two cases.

The overall results are thus consistent with earlier descriptions of the UV-phenotype, which referred to a more “stocky” architecture (Barnes, Ballaré & Caldwell, 1996; Jansen et al., 1998; Robson et al., 2015b). Other aspects of the UV-phenotype, such as leaf thickening are also apparent in the current study, and a decrease in SLA was noted in plants exposed to UV-A enriched radiation. A similar UV induced stocky phenotype has been observed in a substantial number of plant species (Robson et al., 2015b). Yet, this phenotype remains an enigma, in that major questions remain to be answered with respect to the wavelength specificity of its induction, the underlying mechanism of the response, and the functional importance of the induced architectural response for the plant.

### 4.2. The stocky UV phenotype is not associated with plant stress

Several hypotheses have, over the years, been proposed to explain the mechanism underlying UV-induced changes in plant morphology (Robson et al., 2015b). High UV intensities can drive the development of stress induced morphogenic responses (SIMR) as first proposed by Potters, Pasternak, Guisez, Palme & Jansen (2007). These responses are associated with the disruption of cellular metabolism (distress), resulting in a localized cessation of growth. The SIMR phenotype is characterised by decreased elongation growth, which can result in a more stocky phenotype, as described in the current study. However, in the current study there is no evidence for disruptive stress. There was a small increase in NPQ in plants grown under UV-A-enriched light but no change was found in neither maximum photochemical efficiency, redox state of PSII nor the operation efficiency of PSII in any of the UV treatments measured using chlorophyll *a* fluorometry, nor in total chlorophyll content. Furthermore, there is no evidence for an increase in accumulation of damaged DNA, i.e. CPD dimers. DNA damage may potentially result in a stocky phenotype as UV-induced dimerization of DNA can impair DNA replication, and hence impede cell cycle progression, particularly by slowing the G1-to-S phase (Jiang, Wang & Björn, 2011). Indeed, several earlier reports refer to UV-mediated impairment of cell division (Dickson & Caldwell 1978; Wargent, Gegas, Jenkins, Doonan & Paul, 2009). Others (Lake, Field, Davey, Berrling & Lomax, 2009) refer to larger cells in UV exposed plants which can be explained by endoreduplication, resulting in fewer, but bigger cells (Radziejwoski et al., 2011). Plants showing symptoms of UV stress would often have between 50 and 800 CPDs/Mb (Kang et al., 1998; Kalbin et al., 2001; Pescheck, Lohbeck, Roleda & Bilger, 2014). Yet, in the current study levels of CPDs/Mb were one- to two-orders of magnitude less, indicating efficient repair of damaged DNA. Therefore, the observed stocky phenotype is not associated with accumulated DNA damage. The data on the ABA content also are consistent with non-stress conditions. Furthermore, the data show increases in total antioxidant activity (TAC) and flavonol concentrations in leaf adaxial epidermis (LAEFC) and the entire leaf (TUAP). Taken together, these data reveal successful UV acclimation. Thus, healthy plants display the stocky UV-phenotype, implying that the strong morphological response observed in UV-exposed cucumber seedlings is a regulatory adjustment that is part of the UV acclimation processes involving UV-A and/or UV-B photoreceptors.

### 4.3. UV acclimation is accompanied by a decrease in biomass accumulation

Notwithstanding the apparent lack of plant stress under either UV-A or UV-B enriched light, a substantial decrease in produced total biomass was noted for plants exposed to UV-B enriched light. We have interpreted this decrease in biomass production as a secondary consequence of UV-acclimation. UV-B exposed plants with shorter stems (including shorter internodes) and shorter petioles, will condense the same number of leaves in a smaller area, hence increasing the likelihood of self-shading which, in turn, may decrease overall PAR capture, and hence photosynthetic productivity (Barnes et al., 1996). Consistently, UV-A had considerably smaller impacts on both stem and petiole length, and this was associated with a lack of impact on biomass production. UV-B-induced decreases in leaf area will further hamper PAR capture, an effect that is much smaller in plants raised under UV-A enriched light. These UV-B-mediated decreases in light capture, may also be accompanied by the often-reported UV-B-induced stomatal closure (He et al., 2013; Martínez-Lüscher et al., 2013; Tossi, Lamattina, Jenkins & Cassia, 2014), which would similarly decrease photosynthesis *in situ*, and potentially decrease biomass production. As a besides, the measured UV-A-induced decrease in SLA, may potentially improve the water use efficiency of plants (Liu & Stützel, 2004), and although water use efficiency has not been measured in this study, several reports have reported UV-induced co-tolerance with drought (Barnes et al., 2019).

### 4.4. Regulatory mechanism(s) underlying the stocky UV-induced phenotype

Since the data in this study do not support an association between the stocky UV phenotype and plant stress, one or more different specific regulatory response should be considered, as argued above, dependent on what part of the UV spectrum is supplementing the PAR. The UV-B photoreceptor UVR8 was discovered in Arabidopsis mutants that did not show UV-B-induced dwarfing of hypocotyls (Hayes, Velanis, Jenkins & Franklin, 2014). Thus, UVR8 has been strongly associated with control of plant architecture. It should also be noted that, in a recent study by Rai et al. (2020), there was a marked difference in regulation of gene expression between a UVR8-dominated effect at wavelengths below 335-350 nm and a cryptochrome-dominated regulatory mechanism at UV wavelengths above 350 nm. Notwithstanding, the UVR8- and CRY-regulatory mechanisms were interdependently influenced by each other.

Thus, the mechanism underlying the UV-B induced stocky phenotype may relate to interactions with various cellular signalling pathways, including the phytochrome and cryptochrome pathways. In UV-exposed plants, UVR8 monomers bind COP1, and the resulting UVR8-COP1 complexes enter the nucleus and promote UV-B signalling which inhibits auxin biosynthesis, signalling as well as hypocotyl elongation (Hectors, van Oevelen, Guisez, Prinsen & Jansen, 2012). Parallel increases in the expression of the HY5/HYH transcription factor may enhance transcription of polar auxin transport proteins PIN1 and PIN3, as well as several regulators of auxin signalling (Vanhaelewyn, Prinsen, van der Straeten & Vandenbussche, 2016). Furthermore, sequestration of COP1 by UVR8 destabilises PIF5, further interfering with auxin biosynthesis and signalling (Hayes et al., 2014; Vanhaelewyn et al., 2016). Yet, few studies have been able to demonstrate UV induced changes in auxin levels. Hectors et al. (2012) reported (non-significant) UV-B induced decreases in auxin levels in Arabidopsis leaves, as well as altered UV-B responses in auxin influx and biosynthesis mutants (Hectors et al., 2012). The current study does not present evidence for significant changes in auxin concentrations either, despite strong morphological responses, but a non-significant trend of decreasing GA concentrations is observed in both UV-A and UV-B exposed plants. UVR8 binding to COP1 can, via upregulation of transcription of HY5 and HYH, result in an increase in GA2ox1 levels, reducing GA concentrations (Hayes et al., 2014; Vanhaelewyn et al., 2016). The current study shows a trend of decreasing concentrations of GA1, GA44, GA9 and GA15, in both UV-A and UV-B exposed plants. The data in the current paper are not conclusive with respect to a role for UVR8, auxin or gibberellic acid in mediating plant UV-responses. In fact, it is debatable whether the observed dwarfing response can simply be explained as UVR8-mediated, given that responses are induced by both UV-B and UV-A radiation, albeit with partly different outcomes (progressive decreases in petiole lengths and leaf area in plants grown in UV-B-enriched light; alteration in carbon allocation from shoots toward roots in UV-A-enriched light). Therefore, although the UVR8 action spectrum remains to be fully characterised in detail, particularly with respect to potential interactive responses to UV-A wavelengths, recent evidence suggests antagonistic effects whereby responsivity to UV-B is modulated by UV-A wavelengths and *vice versa* (Morales et al., 2013; Rai et al., 2020). Thus, although a UVR8-mediated mechanism appears to be the most likely candidate to explain observed decreases in both organ elongation and GA concentration in UV-B-exposed plants, major questions remain to be addressed concerning the regulation of the stocky phenotype under natural, solar light conditions where there is considerable scope for interactions between multiple wavelength bands, photoreceptors, and signalling pathways. In fact, brassinosteroids may also play a role in UV-regulation of gene expression (Sävenstrand, Brosché & Strid, 2004) and development (Liang et al., 2018), a role which was further strengthened by the discovery of interaction of UVR8 with molecular regulators of brassinosteroids.

### 4.5. Relationships between flavonoid content and antioxidant capacity

Protection against UV was measured in three different ways reflecting the notion that UV-B-induced flavonoidal compounds have both an antioxidative effect and a function as UV-absorbing compounds (Agati, Azzarello, Pollastri, & Tattini, 2012; Hideg & Strid, 2017). For estimation of leaf adaxial epidermal flavonol content (LAEFC), the Dualex method was used, for TUAP acidic methanol extraction and spectrophotometric detection at 330 nm was employed, and total anti-oxidative activity (TAC) was assayed using the Trolox method. Comparisons between LAEFC and TUAP measurements have been made previously (Barthod, Cerovic & Epron, 2007), and in line with published results the intercept of the plot of LAEFC versus TUAP measurements (Supplementary Fig. S1) is inferring higher flavonol content in the epidermal cell layers compared with the average in the total foliar biomass.

The data of this paper show an increased flavonoid level in leaves for up to five days after all three treatments (no UV control, UV-A-, or UV-B-enriched light). Thereafter the flavonoid content declined (Figs 8 A and B). The dependence of the flavonoidal levels (LAEFC and TUAP), with a peak at day 5 is likely to be due to leaf developmental processes. TAC also peaked on day 5 (Fig. 8C). That the peaks in all three parameters at day 5 could be related to particular weather conditions seems highly unlikely given that all experiments were independently replicated three times during different weeks between February and June. For TUAP, readings after 14 days of UV exposure were similar to those measured at the onset of the experiment (day 0). For the LAEFC, the readout on day 14 returned to levels slightly lower than at the outset, whereas for TAC the antioxidant capacity decreased between day 0 and day 14 by about 50%, independently of whether the plants had experienced any UV exposures or were controls. Thus, as these trends were observed in both control and in UV-treated plants, a developmental process would be the likely cause. To improve understanding of developmental processes as well as the relationships between the three different parameters, further regression analyses were performed.

The shape of the TAC curve of antioxidant capacity as compared LAEFC or TUAP results indicates that the TAC parameter reflects two different physiological means of antioxidative activity, one being flavonoids and the second being another type of ROS scavenging of enzymatic or non-enzymatic nature, and which shows an age-dependent decrease by half through days 10 to 14. When estimating the linear dependence of the TAC and LAEFC experiments, we found that the intercept at a zero TAC level was similar for the two leaf-age-dependent linearizations used (Eqs. 2 and 3), with a LAEFC flavonol index of approximately 0.13 (Fig. 9A), which indicates the presence of some flavonoidal compounds that lack antioxidant capacity, e.g. monohydroxylated species (Tattini et al., 2012). Differences in the antioxidant capacity of different flavonol species have been extensively demonstrated (Csepregi, Neugart, Schreiner & Hideg, 2016). For example, some flavones, such as apigenin, have particularly low ferric ion reducing antioxidant power (Csepregi et al., 2016). Also, it should be considered that Trolox may not be a perfect proxy for antioxidant activity in general (Csepregi et al., 2016).

The difference in the slope of the two linear relationships in Supplementary Fig. 9A and the different shapes of the curves in Fig. 8A and 8B indicate that, as the leaf tissue aged, the pool of leaf epidermal flavonols had lost its anti-oxidative capacity, either by accumulation of more oxidized forms of this type of compounds, because of increased O-glycosylation, or due to another yet to be identified mechanism. The OH group on the 3-position on the A-ring of the flavonoid backbone is commonly glycosylated, which decreases antioxidant activity. Developmental changes in flavonol profile have also previously been shown in for instance *Sinapis alba* (Reifenrath & Müller, 2007) and *Vitis vinifera* (Bouderias, Teszlak, Jakab, & Körösi, 2020). Also, a study by Morgenstern, Ekholm, Scheewe & Rumpunen (2014) shows that the ratio of quercetin to kaempferol in buckthorn shows a strong developmental trend, rising from just over 100 at the beginning of the season, to over 400 by mid-summer, and this effect is paralleled by a drop in gallic acid and rutin levels (Morgenstern, et al. 2014). Given the different antioxidant activities of different flavonols, this may underpin a change in antioxidant capacity.

In our study, the loss of anti-oxidative defense (i.e. TAC) was independent of whether the plants had been kept under control conditions or under supplementary UV-A- or UV-B-enriched light. Thus, this seems to be a true effect of leaf age. In old leaves, flavonoids constitute the bulk of the antioxidant capacity (intercept close to 0 in Supplementary Fig. 9B), whereas in younger tissue half of the oxidant capacity (comparison of the curves in Figs 8B and 8C and considering the intercept at 2.37 in Supplementary Fig. 9B) is contributed by antioxidative systems other than flavonoids, be it enzymatic or non-enzymatic, that do not absorb light at 330 nm in methanol under acidic conditions. Leaf age-dependent changes in TAC have previously been shown in greenhouse-grown grapevine leaves (Majer & Hideg, 2012), using a several-fold higher biologically effective UV-B dose than we did. Four days of exposure led to a large increase in TAC in young leaves, similarly to what we found in cucumber. However, in old leaves, 4 days of UV exposure led to decreased TAC. Clearly, there are strong interactions between UV acclimation and developmental processes that govern as disparate physiological parameters in plants as stem and petiole stretching, leaf expansion, flavonoid content and total antioxidant capacity.

### 4.6. Conclusion

In this paper we show that cucumber grown in UV-A- or UV-B-enriched light led to a stockier phenotype compared to on-UV-irradiated control plants. In plants grown in UV-A-enriched light, the decreases in stem and petiole lengths were similar independently of tissue age whereas in plants grown in UV-B-enriched light stems and petioles were progressively shorter the younger the tissue. In addition, plants grown under UV-A-enriched light significantly reallocated photosynthate from shoot to root, had thicker leaves and decreased specific leaf area. This infers different morphological plant regulatory mechanisms under UV-A and UV-B radiation, especially since there was no evidence of stress in any of the UV-exposed plants, as judged by the absence of effects on photosynthetic parameters, cyclobutane pyrimidine dimer levels, or ABA content. The total leaf antioxidant activity and UV-dependent accumulation patterns of flavonoidal compounds and leaf-age-dependent variation in these parameters also indicated successful acclimation of the plants to the two UV light regimens. Therefore, we conclude that the stocky UV phenotype developed in healthy plants, which in turn implies a strong regulatory adjustment and morphological response as part of a successful UV acclimation processes involving UV-A and/or UV-B photoreceptors.

## Acknowledgements

This research was supported by grants to Å.S. from the Knowledge Foundation (kks.se; contract no. 20130164), and the Swedish Research Council Formas (formas.se/en; Contract no. 942-2015-516). The Örebro University’s Faculty for Business, Science and Technology also supported the research. M.A.K.J acknowledges support by Science Foundation Ireland (S16/IA/4418). E.P acknowledges support by the Flemish Science Foundation (FWO, grant G000515N). M.Q. was sponsored by the China Scholarship Council (CSC no. 201406320076).

## Author contribution

Åke Strid, Marcel Jansen, Eva Rosenqvist, Minjie Qian and Irina Kalbina planned the research. Minjie Qian, Åke Strid, Marcel Jansen, Eva Rosenqvist, Els Prinsen, and Frauke Pescheck designed experiments. Minjie Qian, Els Prinsen, Frauke Pescheck and Irina Kalbina performed experiments. Minjie Qian, Els Prinsen, Åke Strid, Frauke Pescheck, Ann-Marie Flygare, Eva Rosenqvist and Marcel Jansen analyzed data. Åke Strid and Marcel Jansen wrote the paper with contributions from Minjie Qian, Eva Rosenqvist, Ann-Marie Flygare, Els Prinsen and Frauke Pescheck. All authors commented and approved the manuscript.

## Supporting information

Additional supporting information may be found online in the Supporting Information section at the end of this article.

