## Supplementary Text+Figs for "Downsizing in plants - UV induces pronounced morphological changes in cucumber in the absence of stress"

**Modelling the relationship between leaf flavonoid content and antioxidant capacity**

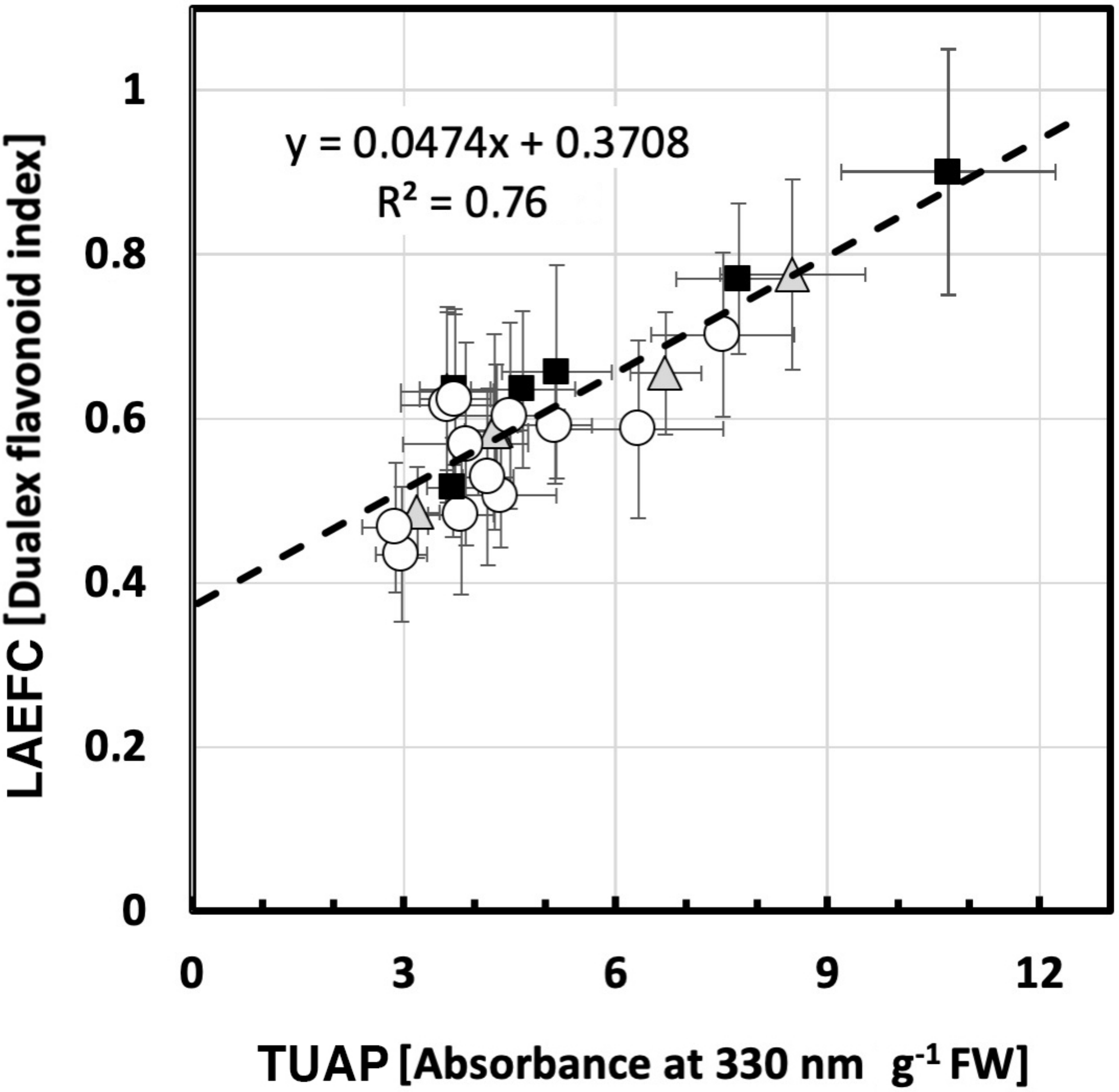


**Supplementary Fig. S1.** The monophasic relationship assumed in Eqn. S1 between LAEFC (Dualex) and TUAP. Open circles denote UV-deficient control treatment, grey triangles denote treatment with UV-A-enriched light and black boxes denote treatment with UV-B-enriched light.

**LAEFC vs. TUAP**

To ascertain to what extent the methods used for LAEFC and TUAP leaf flavonoid content describe the same physiological process within the plant leaf (i.e. that the two different pools of flavonoids measured by these techniques where directly proportional to each other), two models were adapted to describe the relationship between the methods. In Supplementary Figure S1, a simple linear relationship between read-outs of the two analytical methods was assumed, without taking into account the type of treatment (UV-A- or UV-B-enriched) or leaf age. The linear relationship was found to be:

$$\hat{y} = 0.37 + 0.047X, \qquad \text{(Eqn. S1)}$$

where $\hat{y}$ is the estimated expected LAEFC Dualex read-out and x is the value when TUAP method is used. The $R^2$ of this linear fit was 0.76. In the second model, the dependence between the results of the LAEFC and TUAP methods was assumed to be due also to treatment (UV-A- or UV-B-enriched, or control) and leaf age (see main text Eqn. 1). A small influence of leaf age was found with a $R^2$ of this fit was 0.84, indicating that the second model is better in explaining the dependence between LAEFC and TUAP. Also, the residual plots show a random pattern on both sides of 0 (Supplementary Fig. S2A and S2B), thus justifying the model assumptions. Thus, as the tissue aged, there was a tendency for the first model (Supplementary Eqn. S1) to exhibit increased residuals (deviations between true and estimated values), indicating that leaf age is also a determinant of the relationship between LAEFC and TUAP.

**TAC vs. LAEFC**

The dependence between the TAC and the LAEFC assays, was first studied assuming two simple linear relationships, without taking into account the type of treatment (UV-A- or UV-B-enriched), but dividing up the samples in those for younger leaves (≤ 5 days of exposure time, i.e. ≤ 19 days after sowing) and those from older leaves (≥ 10 days of exposure time, i.e. ≥ 24 days after sowing) (see Main Text Eqs. 2 and 3).

We also applied a second model to describe the results of the LAEFC measurements with the TAC method as an explanatory variable and also including explanatory variables treatment (UV-A- or UV-B-enriched) and leaf age. However, in this case we did not find any statistically significant proof for this assumption, and instead obtained a straight line dependence with the following equation:

$$\hat{y} = 0.34 + 0.08X, \qquad \text{(Eqn. S2)}$$

with an $R^2$ value 0.70. The intercept at 0 TAC now gave a value of 0.34 in the LAEFC parameter. The plot of the residuals for the second model is shown in Supplementary Fig. S2C, again exhibiting a random pattern on both sides of 0 indicating that this model is sufficient to explain the data. Thus, we could not conclude with statistical significance whether a leaf age effect was explaining the correlation between the data or not.

### TAC vs. TUAP

The dependence between TAC and TUAP was then studied. In Eqs. 4 and 5 and Fig. 9B of the Main Text, we first assumed two simple linear relationships between TAC and TUAP with $R^2 > 0.91$ in both cases. Also, in the second model, where involvement of both treatment effects and leaf age, were considered. A clear effect of tissue age was seen on the dependence between the TUAP and TAC assays. In this case, the estimated equation became:

$$\hat{y} = 2.07 + 0.39X - 0.13X3, \qquad \text{(Eqn. S3)}$$

again with a high factor $R^2 = 0.96$. The residual plots (Supplementary Figs. S2D and S2E) corroborate this choice of model.

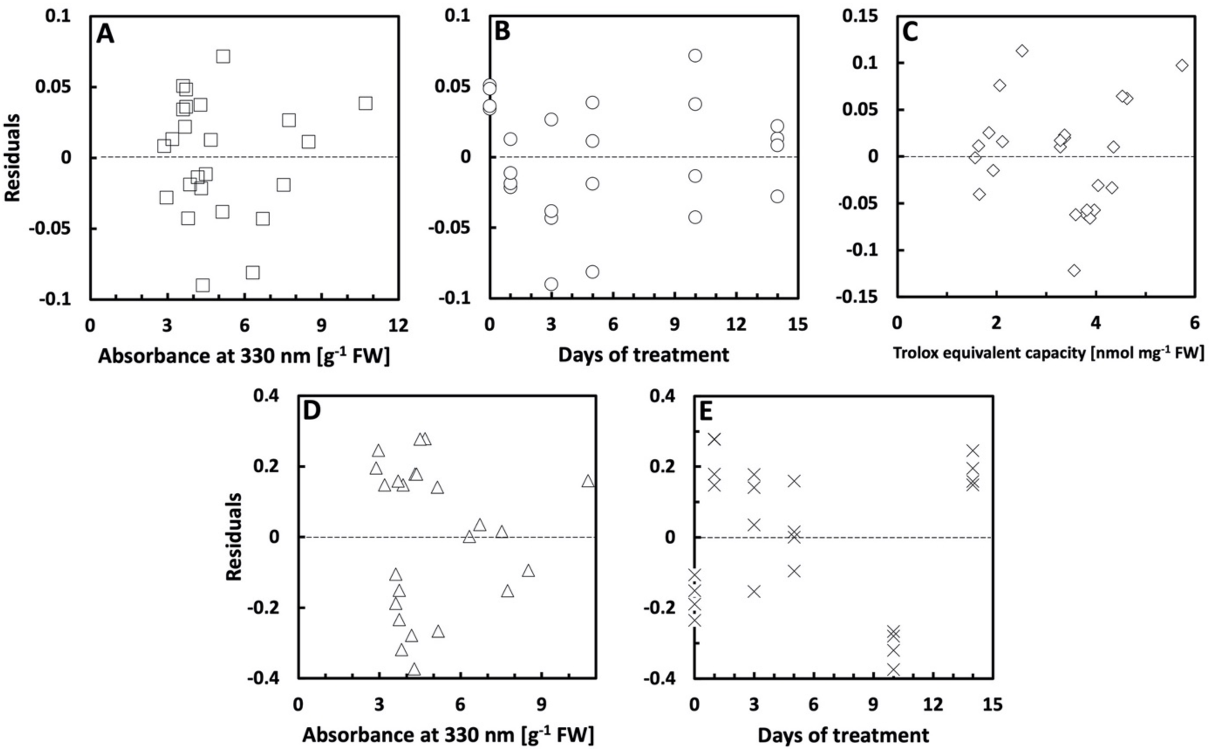


**Supplementary Figure S2.** Residual (ε) plots for models of correlation between the three methods used to determine LAEFC, TUAP, and TAC (see Fig. 9 of the main text and Supplementary Fig. S1). A) Residual plot for the OD330 variable of the correlation of between LAEFC and TUAP; B) Residual plot for the leaf age variable of the correlation between LAEFC and TUAP; and C) Residual plot for the TAC variable of the correlation of between LAEFC and TAC; D) Residual plot for the TUAP variable of the correlation of between TAC and TUAP; E) Residual plot for the leaf age variable of the correlation between TAC and TUAP.
